# PROTAC-mediated selective degradation of cytosolic soluble epoxide hydrolase enhances ER-stress reduction

**DOI:** 10.1101/2022.01.20.477134

**Authors:** Yuxin Wang, Christophe Morisseau, Akihiro Takamura, Debin Wan, Dongyang Li, Simone Sidoli, Jun Yang, Dennis W. Wolan, Bruce D. Hammock, Seiya Kitamura

**Affiliations:** Department of Entomology and Nematology, and UC Davis Comprehensive Cancer Center, University of California Davis, One Shields Avenue, Davis, CA 95616, United States; Department of Molecular Medicine, The Scripps Research Institute, La Jolla, CA 92037 USA; Department of Integrative Structural and Computational Biology, The Scripps Research Institute, La Jolla, CA 92037 USA; Department of Biochemistry, Albert Einstein College of Medicine, Bronx, NY 10461 USA

## Abstract

Soluble epoxide hydrolase (sEH) is a bifunctional enzyme responsible for lipid metabolism and is a promising drug target. Here, we report the first-in-class PROTACs small molecule degraders of sEH. Our optimized PROTAC selectively targets the degradation of cytosolic but not peroxisomal sEH, resulting in exquisite spatiotemporal control. Remarkably, our sEH PROTAC molecule has higher potency in cellular assays compared to the parent sEH inhibitor as measured by significantly reduced ER stress. Interestingly, our mechanistic data indicate that our PROTAC directs degradation of cytosolic sEH via the lysosome, not through the proteasome. The molecules presented here are useful chemical probes to study the biology of sEH with the potential for therapeutic development. Broadly, our results represent a proof-of-concept for the superior cellular potency of sEH degradation over sEH enzymatic inhibition, as well as subcellular compartment-selective modulation of a protein by PROTACs.

**Highlights:**

- First-in-class soluble epoxide hydrolase (sEH) small-molecule degraders.
- Selective degradation of cytosolic but not peroxisomal sEH.
- Significant and stable reduction in sEH protein levels, leading to enhanced cellular efficacy in ER stress reduction relative to the parent inhibitor.

## Introduction

Soluble epoxide hydrolase (sEH) is a bifunctional enzyme present in vertebrates and is encoded by the *Ephx2* gene (Newman, Morisseau et al., 2003). The N-terminal domain possesses a lipid phosphatase activity, while the C-terminus is responsible for metabolizing epoxy fatty acids to corresponding 1,2-diols (Hashimoto, 2019, Morisseau & Hammock, 2013). Inhibition of sEH hydrolase activity by small molecules is a promising approach for the treatment of multiple diseases including inflammation, pain, cancer, and metabolic disorders (Fishbein, Wang et al., 2020, Wagner, McReynolds et al., 2017, Wang, Yang et al., 2018). Notwithstanding the extensive biological knowledge and design of small-molecule inhibitors for the sEH domain, the biological importance of the N-terminal phosphatase domain remains to be elucidated.

Mammalian sEH is localized to both the cytosol and peroxisomes (Gill, Grant et al., 1994). A previous study indicated sEH contributes to stroke injury only when localized in the cytoplasm, while peroxisomal sEH may be protective, showing the different biological functions of cytosolic and peroxisomal sEH (Nelson, Zhang et al., 2015). Interestingly, the human sEH is located in both cytosolic and peroxisomal compartments in hepatocytes and renal proximal tubules, while sEH is exclusively located in cytosolic compartment in other sEH-containing tissues such as pancreatic islet cells, intestinal epithelium, and anterior pituitary cells. These data indicate that sEH subcellular localization is tissue-dependent and sEH may have tissue- and subcellular location-specific functions (Enayetallah, French et al., 2006). Therefore, any molecule that could preferentially label and/or inhibit cytosolic vs. peroxisomal sEH (and vice versa) would be an essential tool to understand the differences in biological importance of sEH between the two cellular locations.

Proteolysis targeting chimera (PROTAC) is a promising technology that promotes the highly selective degradation of target proteins within cells and whole animals (Sakamoto, Kim et al., 2001, Sun, Gao et al., 2019). PROTACs are heterodimeric molecules that recruit an E3 ligase to proteins of interest for ubiquitination and proteasomal dependent degradation (Sun et al., 2019). In comparison to traditional reversible and covalent small-molecule inhibitors, PROTACs have the additional advantage of labeling and degrading those proteins considered “undruggable”, including (but not limited to) transcription factors, transcriptional regulators, and protein-protein interaction partners (Bai, Zhou et al., 2019, Bond, Chu et al., 2020, Garnar-Wortzel, Bishop et al., 2021, Neklesa, Snyder et al., 2019). Conversion of established and modest small-molecule protein binders into PROTACs has resulted in improved cellular selectivity and potency. This has significant implications for therapeutics because lower PROTAC concentrations are required for cellular activity relative to the traditional therapeutic counterpart (Salami, Alabi et al., 2018, Zou, Ma et al., 2019). Additionally, PROTACs are “catalytic” and deviate significantly from the mode of actions of traditional competitive- and occupancy-driven inhibitors (Gao, Sun et al., 2020). Altogether, small-molecule degraders, including PROTACs, are an attractive modality in drug development to overcome the limitations of traditional occupancy-driven ligands.

Here, we report the development of the first-in-class small-molecule degraders of sEH. We demonstrate their ability to reduce sEH levels in a spatial selective manner. We further showed enhanced cellular potency of the PROTAC in the cellular ER stress assay compared to the parent sEH inhibitor. Finally, our mechanistic studies indicate that the degradation mechanism of sEH induced by our PROTAC is lysosome-dependent and not proteasome-dependent, contrary to the current assumption of degradation mechanism induced by PROTACs.

## Materials and Methods

### Cell Culture and reagents

Human embryonic kidney 293T cells and human hepatocyte carcinoma HepG_2_ cells were purchased from American Type Culture Collection (ATCC, Manassas, VA), and the ATG 2A/2B ^-/-^ cell line was a kind gift from Dr. Komatsu Masaaki (Juntendo University, Tokyo, Japan). The cell lines were cultured in high-glucose (4.5 g/L) Dulbecco’s Modified Eagle’s Medium (HyClone, Marlborough, MA). The media was supplemented with 10% fetal bovine serum (VWR, Radnor, PA), 100 units/mL of penicillin, and 100 mg/mL of streptomycin (Thermo Fisher Scientific, Waltham, MA). The cells were cultured at 37 °C in a 5% CO_2_ incubator. Cycloheximide (CHX, #C1988), MG132 (#474790), VER155008 (SML0271) were purchased from Sigma-Aldrich (St. Louis, MO), Bortezomib (BTZ, #NC0175953) was purchased from LC laboratory (Woburn, MA), bafilomycin A1 (BafA1, #54645S) was purchased from Cell Signaling Technology (Danvers, MA), the thapsigargin (Tg, #1138) was purchased from R & D SYSTEMS INC (Minneapolis, MN), and E3 ligase recruiter building blocks thalidomide - linker 2 (#6468/25) were purchased from Tocris Bioscience (Minneapolis, MN) and VHL Ligand 1(#21591) was purchased from Caymen Chemical (Ann Arbor, MI). The detailed synthetic procedure is described in the supporting information.

### Measurement of biochemical inhibitory potency of PROTAC compounds against human and mouse sEH-hydrolase

The inhibitory potency of compounds was measured using a fluorescent assay as previously described (Jones, Wolf et al., 2005). The IC_50_ values are defined as inhibitor concentrations that reduced the enzymatic activity by 50% and were calculated in the linear range of the dose-response curve for log inhibitor concentration versus % inhibition. These values are the averages of three replicates, obtained from the experiments with various concentrations of the inhibitors (1–10,000 nM) including at least two datum points above and two datum points below the IC_50_. We previously showed a high z-value for the assay and the standard error for the IC_50_ values is between 10 and 20% for the assays performed here suggesting that the differences of two-fold or greater would be significant.

### Measurement of phosphatase activity of sEH

The soluble epoxide hydrolase phosphatase activity was determined by quantitative analysis of 1-myristoyl-glycerol (product) using LC-MS/MS as previously described (Morisseau, Schebb et al., 2012). Briefly, one μL of a 5 mM solution of 1-myristoyl-glycerol-3-phosphate in water ([S]_final_ = 50 μM) was added to 100 μL of cell extracts. The reaction mixtures were incubated at 37°C for 5 to 30 minutes. The reactions were then quenched by adding 100 μL of a 50:49:1 mixture of acetonitrile, water and acetic acid containing 200 nM of hexanoyl-ceramide as internal standard. The chromatographic analysis was carried out on a Supelco Discovery “Bio Wide Pore” C-5 RP pre-column (Sigma–Aldrich, St Louis, MO) and the mass spectrometer was operated in scheduled MRM mode with optimized MS transitions as previously described (Morisseau et al., 2012).

### Measurement of inhibitory potency against human mEH

The inhibitory activity of the PROTAC compounds toward human microsomal epoxide hydrolase (mEH) was determined by a fluorescence-based assay, using purified recombinant human mEH protein and cyano(6-methoxy-naphthalene-2-yl) methyl glycidyl carbonate (CMNG) as a fluorescent substrate (Morisseau, Bernay et al., 2011). The enzymes were incubated at 37 °C with the compounds ([I]_final_ = 400-100,000 nM) for 5 min in 100 mM Tris/HCl buffer (200 μL, pH 8.5) containing 0.1−1 mg/mL of BSA and 1% of DMSO. The substrate (CMNG) was then added ([S]_final_ = 5 μM). The activity was assessed by measuring the appearance of the fluorescent 6-methoxynaphthaldehyde product (λ_ex_ = 330 nm, λ_em_ = 465 nm) every 30 s for 10 min at 37 °C on a SpectraMax M2 microplate reader (Molecular Devices). All measurements were performed in triplicate and the mean was reported.

### Compound treatment and immunoblotting analysis

Human embryonic kidney (HEK) 293T cells and human hepatocyte carcinoma HepG_2_ cells were treated with sEH PROTAC **1a**, DMSO vehicle or negative control **1a’** in complete medium. After 24 hours of incubation, the medium was decanted. Then the cells were washed with cold phosphate-buffered saline (PBS) and lysed using RIPA buffer with protease inhibitor cocktail (#5871S, Cell Signaling, Danvers, MA) and phosphatase inhibitor cocktail (#78420, Life Technologies, Carlsbad, CA). The cell lysates were then resolved using SDS/PAGE (Bio-Rad, Hercules, CA) and transferred onto a nitrocellulose membrane using the Trans-Blot Turbo Transfer System (#170-4155, Bio-Rad, Hercules, CA). The membranes were blocked in 5% fat free milk in PBST for 1 hour at room temperature and probed with the primary antibody. The following antibodies were used, including: 1) a previously generated sEH antibody (Imig, Zhao et al., 2002); 2) phospha-IRE1α (#ab48187, Abcam, Waltham, MA); 3) IRE1α (#3294, Cell Signaling, Danvers, MA); 4) XBP-1s (#12782, Cell Signaling, Danvers, MA); and 4) β-actin (#A1978, Sigma-Aldrich, St. Louis, MO). The membranes were then probed with goat anti-rabbit and goat anti-mouse secondary antibodies (#7074, #7076, Cell Signaling, Danvers, MA), and blots were incubated with Clarity Western ECL Substrate (Cat. # 170-5061, Bio-Rad, Hercules, CA) and imaged using the ChemiDoc MP (Cat. # 170-8280, Bio-Rad, Hercules, CA) with ImageLab Version 5 (Bio-Rad, Hercules, CA). Each experiment has three independent repeats.

### Measurement of soluble epoxide hydrolase (sEH) enzyme activity of cell culture

The HepG_2_ cells were treated with different sEH PROTAC compounds and isolated by centrifugation. The cells were suspended in 1 mL of media. To measure the residual soluble epoxide hydrolase activity [^3^H]-*trans*-diphenyl-propene oxide (*t*-DPPO) was used as a substrate (Borhan, Mebrahtu et al., 1995). One microliter of a 5 mM solution of *t*-DPPO in DMSO was added to 100 μL of diluted cell suspension ([S]_final_ = 50 μM). The mixture was incubated at 37 °C for 120 min, and the reaction quenched by addition of 60 μL of methanol and 200 μL of isooctane, which extracts the remaining epoxide from the aqueous phase. Parallel extractions of the stopped reaction with 1-hexanol were performed to assess the possible presence of glutathione transferase activity which could also transform the substrate into a more water soluble glutathione conjugate and give a false positive (Borhan et al., 1995); however, none was observed. The enzyme activity was followed by measuring the quantity of radioactive diol formed in the aqueous phase using a scintillation counter (TriCarb 2810 TR, Perkin Elmer, Shelton, CT). Assays were performed in triplicate. Protein concentration was quantified using the Pierce BCA assay (Pierce, Rockford, IL), using fraction V bovine serum albumin (BSA) as a calibrating standard.

### Quantitation of sEH protein level using a PolyHRP ELISA

The sEH level in HepG_2_ cells was measured using an ultrasensitive PolyHRP based immunoassay, as previously described (Li, Cui et al., 2017). The microplate (No. 442404, Nunc Cat. Thermo Fisher Scientific, Waltham, MA) was coated with anti-human sEH rabbit serum (1:2000 dilution) in 0.05 M pH 9.6 carbonate-bicarbonate buffer overnight at 4 °C. Then the plate was blocked with 3% (w/v) skim milk in PBS for 1 h at room temperature. Human sEH standards and samples with different dilutions in PBS containing 0.1 mg/mL bovine serum albumin (BSA) were then applied in the wells. The biotinylated sEH nanobody (1 μg/mL) in PBS was added together to each well immediately to proceed with the immunoreaction for 1h at room temperature. Then, the SA-PolyHRP in PBS (25 ng/mL) was applied to continue the reaction for another 30 min after washing. The 3,3′,5,5′-tetramethylbenzidine (TMB) substrate was added and incubated for 10-15 min at room temperature. The reaction was stopped using the color development with 2 M sulfuric acid, and the optical density was quantified using a SpectraMax M2 microplate reader (Molecular Devices, Sunnyvale, CA, USA) at 450 nm within 10 min.

### Sub-cellular fractions separation

Sub-cellular fractions were prepared by differential centrifugation as described (Gill & Hammock, 1980). Briefly, the cells were treated with DMSO, **1a’**, or PROTAC **1a** for 24 h, then the cells were homogenized in sodium phosphate buffer (20 mM pH 7.4) containing 5 mM EDTA, 1 mM DTT, 1 mM PMSF. The cells were centrifuged at 100,000 g for 20 minutes at 4 °C. The supernatant was collected as S10 fraction, which contains cytosol and microsome. The pellet was resuspended with the same buffer as described above and further sonicated, then centrifuged at 120,000 g for 12 minutes at 4 °C. The supernatant was collected as P10 fraction, which contains nuclear, mitochondria, and peroxisome.

### Immunofluorescence imaging of cell culture

Cells were seeded in the 96-well plate and treated with DMSO, **1a’**, or PROTAC **1a** for 24 h. The cells were fixed with 4% paraformaldehyde for 20 min and cells were permeabilized using 0.02% Triton X-100 in PBS. After this, the cells were blocked with 5% goat serum, and 1% bovine serum albumin in PBS for 1 hour at room temperature. The cells were then incubated with specific primary antibodies at 4°C overnight and were subsequently incubated with the corresponding secondary antibodies for 1 h at room temperature. The nuclei were counterstained with DAPI for 10 min. Images were acquired at 20x magnification using the ImageXpress Micro XL high-content imaging system (Molecular Devices, Sunnyvale, CA), and automated image analysis was performed using Custom Module Editor, MetaXpress software (Molecular Devices, version 6.2, RRID:SCR_016654). The following primary antibodies were used in this experiment: mouse monoclonal anti-sEH (sc-166961, Santa Cruz Biotechnology, Dallas, TX), LAMP2 rabbit mAb (#49067, Cell signaling, Danvers, MA), β-tubulin antibody (#2146, Cell signaling, Danvers, MA), Catalase rabbit mAb (#12980, Cell signaling, Danvers, MA). The following secondary antibodies were purchased from Invitrogen: Alexa-Fluor 647 (1:500 dilution, Thermo Fisher, Waltham, MA), goat anti-rabbit conjugated to Alexa-Fluor 488 (1:500 dilution, Thermo Fisher, Waltham, MA).

### Proteomics analysis

HepG_2_ cells were treated with sEH PROTAC **1a**, DMSO vehicle or negative control **1a’** in complete medium. After 24 hours of incubation, the medium was decanted, cells were collected, and the cytosol fraction was separated as described above. The protein concentration was measured using BCA Protein Assay Kit (Thermo Scientific). Prior to mass spectrometry analysis, samples were desalted using a 96-well plate filter (Orochem) packed with 1 mg of Oasis HLB C-18 resin (Waters). Briefly, the samples were resuspended in 100 μL of 0.1% TFA and loaded onto the HLB resin, which was previously equilibrated using 100 μL of the same buffer. After washing with 100 μL of 0.1% TFA, the samples were eluted with a buffer containing 70 μL of 60% acetonitrile and 0.1% TFA and then dried in a vacuum centrifuge. For the LC-MS/MS acquisition, samples were resuspended in 10 μL of 0.1% TFA and loaded onto a Dionex RSLC Ultimate 300 (Thermo Scientific), coupled online with an Orbitrap Fusion Lumos (Thermo Scientific). Chromatographic separation was performed with a two-column system, consisting of a C-18 trap cartridge (300 μm ID, 5 mm length) and a picofrit analytical column (75 μm ID, 25 cm length) packed in-house with reversed-phase Repro-Sil Pur C18-AQ 3 μm resin. To analyze the proteome, peptides were separated using a 180 min gradient from 4-30% buffer B (buffer A: 0.1% formic acid, buffer B: 80% acetonitrile + 0.1% formic acid) at a flow rate of 300 nL/min. The mass spectrometer was set to acquire spectra in a data-dependent acquisition (DDA) mode. The full MS scan was set to 300-1200 m/z in the orbitrap with a resolution of 120,000 (at 200 m/z) and an AGC target of 5×10e5. MS/MS was performed in the ion trap using the top speed mode (2 secs), an AGC target of 1×10e4 and an HCD collision energy of 35.

Proteome raw files were searched using Proteome Discoverer software (v2.4, Thermo Scientific) using SEQUEST search engine and the SwissProt human database. The search for total proteome included variable modification of N-terminal acetylation, and fixed modification of carbamidomethyl cysteine. Trypsin was specified as the digestive enzyme with up to 2 missed cleavages allowed. Mass tolerance was set to 10 ppm for precursor ions and 0.2 Da for product ions. Peptide and protein false discovery rate was set to 1%. Following the search, data was processed as described previously (Aguilan, Kulej et al., 2020). Proteins were log2 transformed, normalized by the average value of each sample and missing values were imputed using a normal distribution 2 standard deviations lower than the mean. Statistical regulation was assessed using heteroscedastic T-test (if *P* value < 0.05). Data distribution was assumed to be normal but this was not formally tested.

### Data Analysis

Descriptive statistics were obtained and include mean and standard error of mean for continuous variables and count and proportion for categorical variables. Pairwise comparisons in changes in protein levels and ER stress were performed and include a Student’s t-test for continuous variables and Mann-Whitney test for categorical variables. Comparisons across all groups were made via an ANOVA. The details of statistical analyses are given in the figure legend and supporting information.

## Results

### Development of sEH PROTACs

Two series of sEH PROTAC compounds were synthesized based on sEH inhibitors *trans*-4-[4-(3-trifluoromethoxyphenyl-1-ureido)-cyclohexyloxy]-benzoic acid (*t*-TUCB **1**, series **1**) (Hwang, Tsai et al., 2007) and 12-(3-adamantan-1-yl-ureido) dodecanoic acid (AUDA **2**, series **2**) (Morisseau, Goodrow et al., 1999). Recruiters of E3 ligases (Von Hippel-Lindau (VHL) or cereblon (CRBN)) were connected to sEH binders with various linkers (**Table 1**). Negative control compounds (**1a’** and **2a’, Table 1**) were designed based on compounds **1a** and **2a**. These molecules are expected to have equivalent sEH inhibitory potency compared to the corresponding PROTAC molecules but do not recruit CRBN and do not induce degradation of the target protein.

**Table 1.**
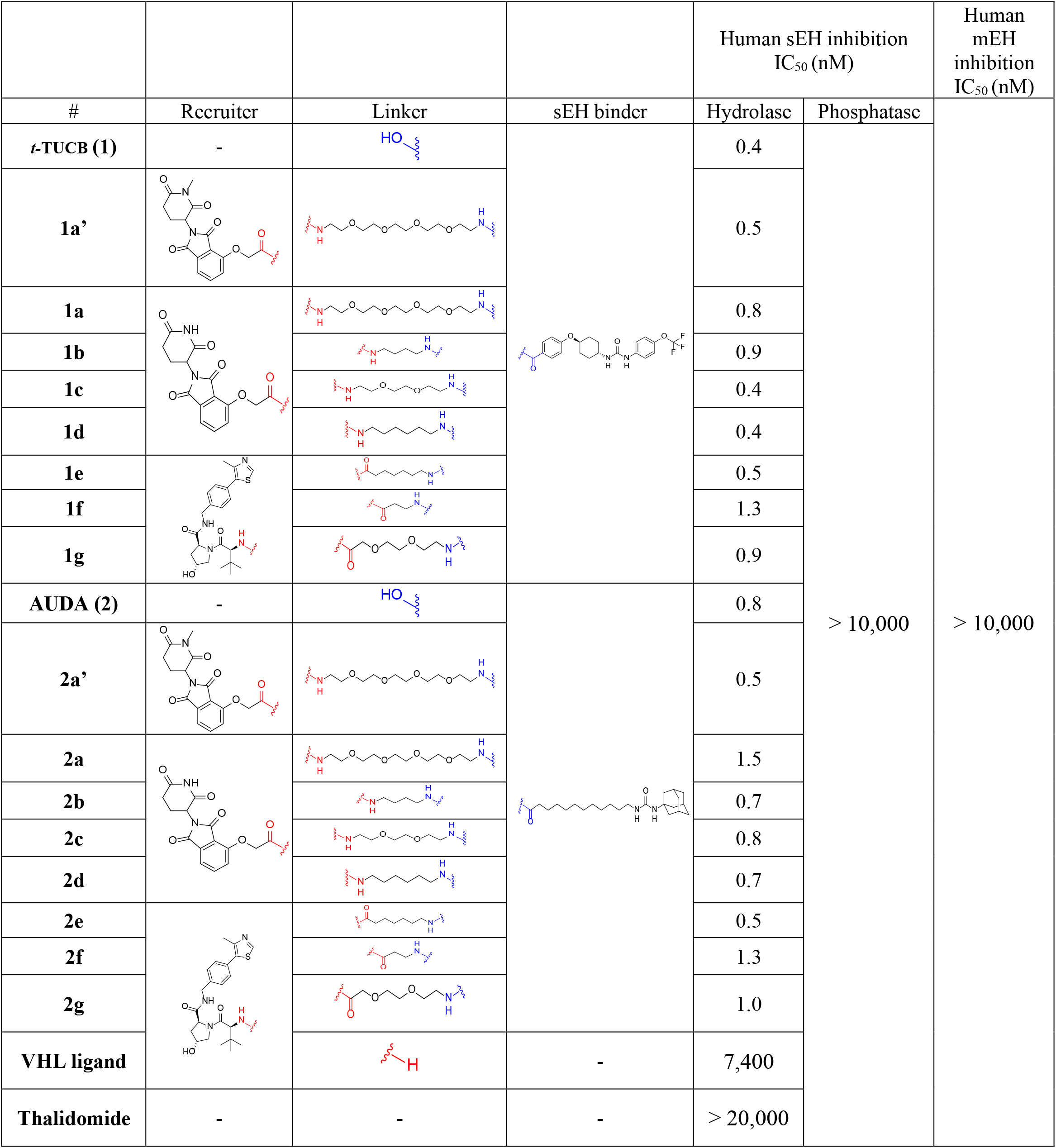
Structure of sEH PROTAC molecules and their inhibitory potency against sEH.

| # | Recruiter | Linker | sEH binder | Human sEH inhibition IC<sub>50</sub> (nM) |  | Human mEH inhibition IC<sub>50</sub> (nM) |
| --- | --- | --- | --- | --- | --- | --- |
|  |  |  |  | Hydrolase | Phosphatase |  |
| *t*-TUCB (1) | - |  |  | 0.4 | > 10,000 | > 10,000 |
| 1a' |  |  |  | 0.5 |  |  |
| 1a |  |  |  | 0.8 |  |  |
| 1b |  |  |  | 0.9 |  |  |
| 1c |  |  |  | 0.4 |  |  |
| 1d |  |  |  | 0.4 |  |  |
| 1e |  |  |  | 0.5 |  |  |
| 1f |  |  |  | 1.3 |  |  |
| 1g |  |  |  | 0.9 |  |  |
| AUDA (2) | - |  |  | 0.8 |  |  |
| 2a' |  |  |  | 0.5 |  |  |
| 2a |  |  |  | 1.5 |  |  |
| 2b |  |  |  | 0.7 |  |  |
| 2c |  |  |  | 0.8 |  |  |
| 2d |  |  |  | 0.7 |  |  |
| 2e |  |  |  | 0.5 |  |  |
| 2f |  |  |  | 1.3 |  |  |
| 2g |  |  |  | 1.0 |  |  |
| VHL ligand |  |  | - | 7,400 |  |  |
| Thalidomide | - | - | - | > 20,000 |  |  |

First, the inhibitory potency against the human and mouse sEH, and human mEH was determined (**Table 1** and **Table S1**). All the molecules showed low to sub nanomolar potency against sEH hydrolase activity, suggesting that the addition of the linker and PROTAC recruiter does not significantly affect compound binding. This is consistent with previous studies showing that a central urea or amide with two lipophilic substitutions is the minimal structural requirement for the sEH inhibition (Imig & Hammock, 2009, Kitamura, Morisseau et al., 2017, Kitamura, Morisseau et al., 2015, Morisseau et al., 1999). Because these molecules are based on the inhibitors of sEH hydrolase, none of them showed inhibition against sEH phosphatase activity nor mEH activity as expected (**Table 1** and **Table S1**).

Encouraged by these biochemical data, next, the sEH protein degradation potency was examined in a cell-based assay using four complementary methods of sEH detection. The human hepatocyte carcinoma cell line HepG_2_ was treated with 1 μM sEH PROTAC compounds for 24 h, and the degradation of sEH was first monitored using immunoblotting. PROTAC compounds **1a, 1c, 1e, 2a, 2c, 2d, 2e** showed degradation of sEH in this assay (**Figure 1A**). There is a correlation of the degradation potency of the molecules with the same linker/E3 recruiter between series **1** and series **2**. For example, a thalidomide recruiter with PEG4 linker (**1a, 2a**) seems to show the highest degradation in each series. These seven compounds (**1a, 1c, 1e, 2a, 2c, 2d, 2e**) were chosen for further analysis. Enzymatic activities were next used to monitor the reduction of the levels of functional sEH (**Figure 1B** and **Table S2**). The parent sEH inhibitors **1** and **2** resulted in only slight inhibition of the hydrolase activity. These compounds are reversible inhibitors and were probably eliminated during the washing step of the cells with fresh media. The low inhibition observed is most likely resulting from the residual inhibitor which remained in the buffer, on the plastic wells, or on the enzyme active site. Consistent with the immunoblotting data, compounds **1a** and **2a** yielded the greatest loss of both hydrolase and phosphatase activity. Finally, we used enzyme-linked immunosorbent assay (ELISA), which can quantitatively determine the absolute concentration of sEH protein in a high-throughput manner (Li et al., 2017). Consistent with other methods, compounds **1a** and **2a** showed the lowest level of sEH protein, suggesting the highest degradation potency (**Figure 1B** bottom panel and **Table S2**). Overall, compound **1a** consistently showed high degradation efficacy in all assays, thus this molecule was selected for further study. To determine the degradation potency, we performed a dose-response (5-1,000 nM) of sEH PROTAC **1a, 1e**, and **2a** in HepG_2_ cells (**Figures 1C** and **S1**), showing that all of them have similar degradation potency in this condition with maximum degradation around 250 nM. A significantly lower amount of sEH was observed at concentrations as low as 25 nM of compound **1a** treatment (**Figure 1C**). As the dose was increased, the sEH protein band became lighter to reach a maximal degradation at 250 nM, then slightly less degradation at 500 nM and 1 μM. The U-shaped concentration-response curve, also known as the hook effect, is a well-known phenomenon observed previously with other PROTACs (Pettersson & Crews, 2019). Interestingly, the treatment failed to reach full elimination of the sEH protein band, even at the maximum effective dose. The degradation efficacy of **1a** at 250 nM was also determined in 293T cells, showing that the compound is effective in both cell lines (**Figure 1D**). We further showed that the co-treatment of sEH inhibitor **1** or **1a’** can counteract the degradation effect induced by **1a**, suggesting that the sEH binding of **1a** is required for the degradation effect of PROTAC (**Figure S2**). In addition, neddylation inhibitor MLN4924 treatment blocked the sEH degradation induced by **1a**, demonstrating that sEH degradation is CRBN-cul-E3 ligase dependent (**Figure S3**). Finally, to demonstrate the selectivity of **1a**, we performed the global proteomics analysis after the treatment of our molecules (**Figure 1E**). The results showed that sEH is one of the most significantly reduced proteins after the treatment of **1a**.

**Figure 1.**
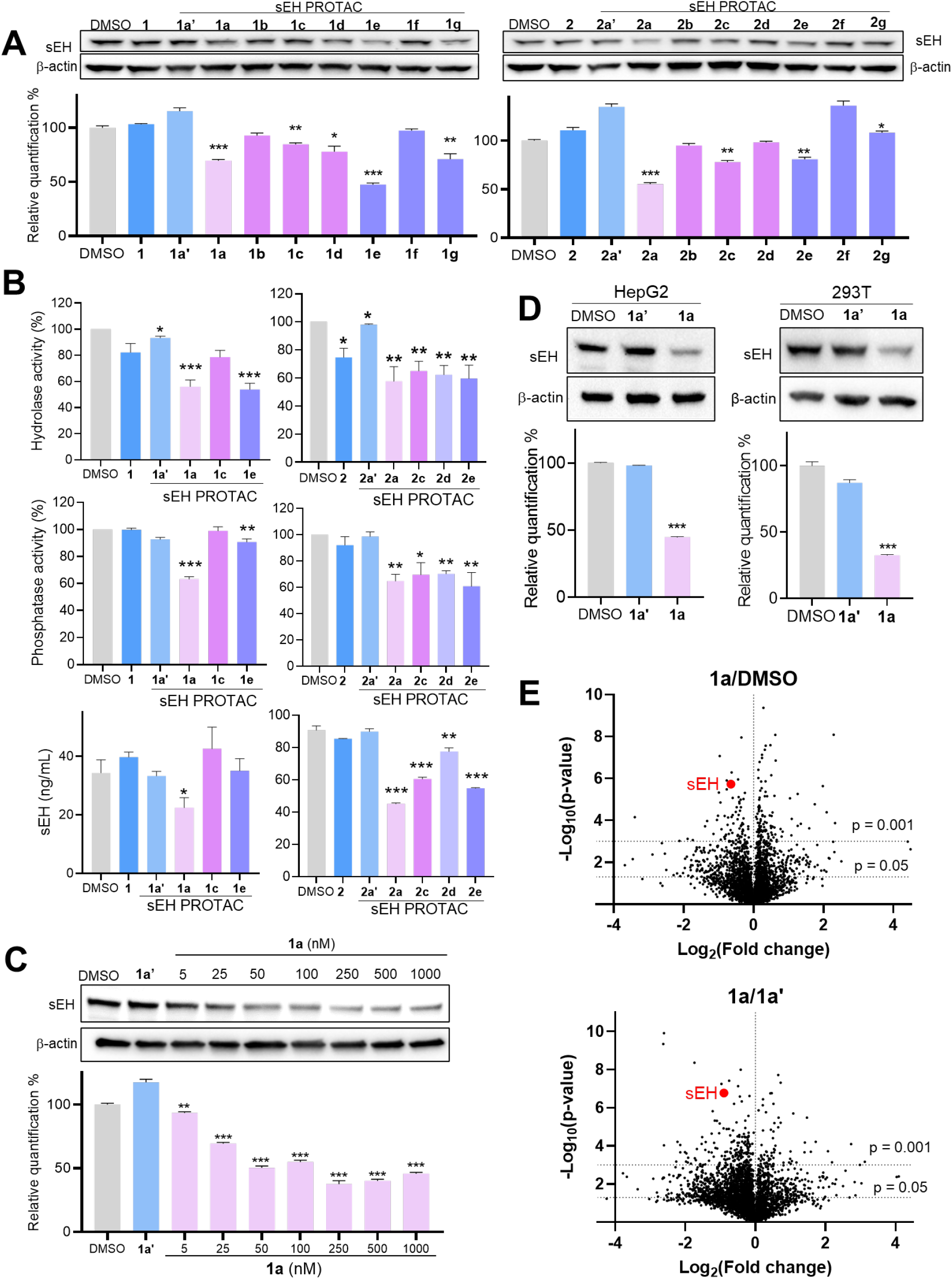
Effect of PROTAC compounds on sEH levels. (**A**) HepG_2_ cells were treated with indicated compound at 1 μM for 24 hours, and the sEH level was measured by immunoblot. Top panel, the representative results of immunoblot; Bottom panel, the quantification of immunoblot results. The structures of sEH PROTAC compounds are shown in **Table 1**. Mean ± SEM are shown. The difference between treatment groups were analyzed by one-way ANOVA with Holm Sidak test, and the normality was analyzed by Shapiro-Wilk test. If there is no significant difference using one-way ANOVA, Student’s t-test was performed. The asterisks in figures **1A, 1B, 1C**, and **1D** represent the significant difference between the compound treatment vs DMSO treatment (*: *P* < 0.05, **: *P* < 0.01, ***: *P* < 0.001). (**B**) HepG_2_ cells were treated with indicated compound at 1 μM for 24 hours, and sEH level was measured by; hydrolase activity (top), phosphatase activity (middle), and ELISA (bottom). All of the *P* values and other statistical details were shown in **Table S2**. (**C**) The concentration response of **1a**. HepG_2_ cells were treated with **1a** for 24 hours and sEH level was measured by immunoblot. Student’s t-test was performed between compound vs DMSO treatment. (**D**) Compound **1a** is effective in both HepG_2_ and 293T cells. HepG_2_ or 293T cells were treated with DMSO, **1a’**, or **1a** (250 nM) for 24 hours, and sEH level was measured by immunoblot. Top panel, the representative results of immunoblot; Bottom panel, the quantification of immunoblot results. The difference between treatment groups were analyzed by one-way ANOVA with Holm Sidak test, and the normality was analyzed by Shapiro-Wilk test. (**E**) Quantitative MS-based proteomic analysis indicates that sEH is one of the most significantly reduced proteins after the treatment of **1a**. HepG_2_ cells were treated with DMSO, **1a**, or **1a’** (250 nM) for 24 hours and proteomic analysis was performed as detailed in the Materials and Methods. The data points representing sEH are labeled in red.

### PROTAC 1a selectively degrades cytosolic sEH but not peroxisome sEH

To explain the apparent lack of total degradation efficacy and given that sEH has a dual subcellular localization (Enayetallah et al., 2006, Moody, Loury et al., 1985), we hypothesized that **1a** induced the degradation of only cytosolic sEH but not the peroxisomal sEH. To test this, the cell lysates were separated into a cytosol-containing fraction (S10) and peroxisome containing fraction (P10). The immunoblot analysis of both fractions showed that **1a** degraded cytosolic sEH while the peroxisomal sEH was not degraded (**Figure 2A**). The hydrolase activity and ELISA results confirmed that compound **1a** selectively degraded the cytosolic sEH (**Figures 2B** and **2C**). Consistent with this, immunofluorescence imaging showed that compound **1a** decreased the colocalization between sEH and the cytosol marker β-tubulin but did not affect the sEH colocalization with the peroxisome marker catalase, further supporting the spatial selectivity of the sEH PROTAC. (**Figures 2D** and **2E**).

**Figure 2.**
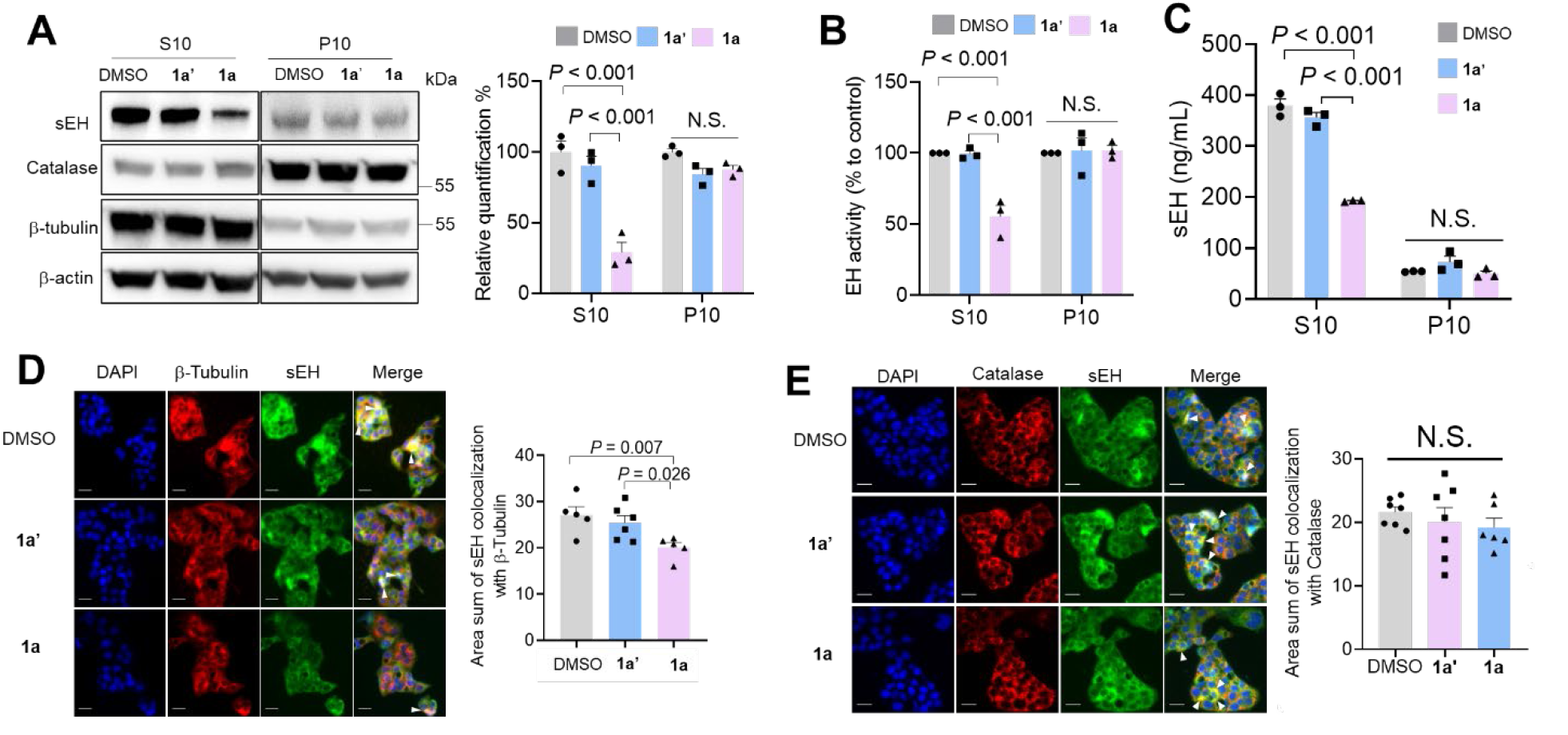
Compound 1a selectively degrades cytosolic sEH but not peroxisomal sEH. HepG_2_ cells were treated with indicated compound at 250 nM for 24 hours and sEH levels in cytosol and peroxisome fractions were measured by; (**A**) Immunoblot. Left panel, the representative results of immunoblot; Right panel, the quantification of immunoblot results. (**B**) Epoxide hydrolase activity, and (**C**) ELISA. (**D, E**) Immunofluorescence results of colocalization of sEH with β-Tubulin (**D**) or catalase (**E**). Scale bar, 25 mm. Left panel, the representative pictures of immunofluorescence, the white arrows indicate the representative merged staining areas of sEH and other proteins; Right panel, the quantification of immunofluorescence results. Mean ± SEM are shown. The difference between DMSO, **1a**’, and **1a** was analyzed using one-way ANOVA Holm Sidak test.

### Probing the kinetics of degradation and production of sEH using PROTACs

Next, the kinetics of sEH degradation induced by **1a** was measured to probe the cellular turnover of sEH. Surprisingly, compound **1a** required 24 h to yield the highest level of degradation (**Figure 3A** and **Figure S4**), unlike previously reported PROTACs that show degradation within 30 min (Liu, Chen et al., 2020). For comparison, the endogenous degradation kinetics of sEH was analyzed using cycloheximide (CHX), the protein synthesis inhibitor (Schneider-Poetsch, Ju et al., 2010). The results showed that it requires 48 h to degrade half of the sEH in the cellular system (**Figure 3B**), indicating that, even slow, PROTAC compound **1a** accelerates the degradation of sEH. In addition, there was no significant difference between CHX treatment with/without the proteasome inhibitor bortezomib (BTZ), suggesting that the endogenous sEH is not degraded through proteasome (**Figure S5**).

**Figure 3.**
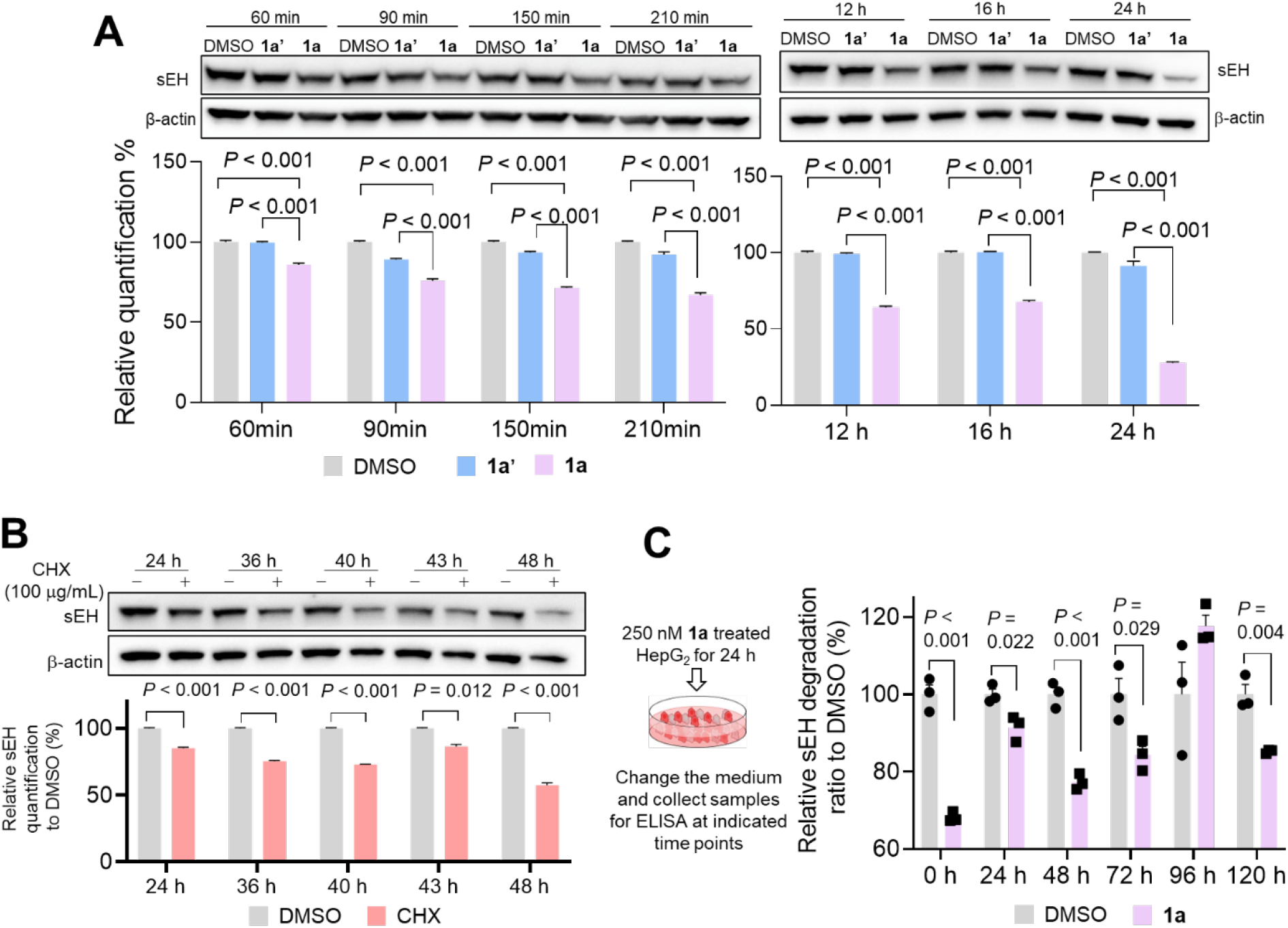
Degradation and production kinetics of sEH. (**A**) Degradation kinetics of sEH induced by PROTAC **1a**. HepG_2_ cells were treated with indicated compound at 250 nM and sEH level was determined by immunoblot. Top panel, the representative results of immunoblot; Bottom panel, the quantification of immunoblot results. Mean ± SEM are shown. The difference between DMSO, **1a**’, and **1a** was analyzed using one-way ANOVA Holm Sidak test. (**B**) Endogenous degradation kinetics of sEH. HepG_2_ cells were treated with protein synthesis inhibitor CHX (100 mg/mL) and sEH level was measured using immunoblot at indicated time points. Top panel, the representative results of immunoblot; Bottom panel, the quantification of immunoblot results. Student’s t-test was performed, and *P* values represent the significant difference between CHX vs vehicle control treatment. (**C**) Probing endogenous sEH production kinetics using PROTAC **1a**. HepG_2_ cells were treated with **1a** at 250 nM for 24 hours, then **1a** was removed from the media, and the sEH protein level was measured using immunoblot after incubating for indicated time. Left panel: the scheme of experiment; Right panel: The quantification of sEH degradation induced by **1a**. Mean ± SEM are shown. Student’s t-test or Wilcoxon-Mann-Whitney test was performed depending on the normality.

We next investigated the kinetics of sEH production using our PROTAC molecule as a chemical probe. HepG_2_ cells were treated with **1a**, then the compound was removed by changing the medium. Cells were incubated in media without PROTAC molecules and collected at different time points. The sEH levels were monitored using the ELISA. The results showed that treatment with **1a** reduced the level of sEH protein up to 72 hours (**Figure 3C**). Altogether, these data show the slow turnover of sEH in the cells and give insights into the endogenous degradation mechanisms and kinetics of sEH.

### Compound 1a appears to degrade sEH through lysosome dependent pathway

PROTACs are believed to trigger the proteasome-dependent degradation of target proteins (Lai & Crews, 2017), thus proteasome inhibitors should be able to rescue the target protein from the degradation effects. To test the mechanism of protein degradation, HepG_2_ cells were treated with proteasome inhibitors BTZ or MG132 in addition to PROTAC **1a**. Interestingly, MG132 failed to block the degradation induced by PROTAC **1a** (**Figures 4A**), and lower concentration of BTZ slightly reversed the sEH degradation induced by **1a** but higher concentrations failed to block the **1a** induced sEH degradation (**Figure S6**), indicating that sEH degradation induced by **1a** is not through the proteasome. In addition, BTZ failed to block the degradation induced by PROTAC molecules **1e** and **2a** that have different sEH binder and PROTAC linker (**Figure S6**), suggesting that all of these structurally diverse molecules induce sEH degradation through proteasome-independent mechanism. Rather, MG132 promoted endogenous sEH degradation (**Figure 4A**). Several studies have reported compensation mechanism between proteasome and autophagy to maintain cellular homeostasis (Ji & Kwon, 2017). Taken together with the slow kinetics, we speculated that sEH is degraded through the lysosomal pathway. To test this possibility, the cells were co-treated with the autophagolysosome inhibitor bafilomycin A1 (BafA1). BafA1 blocked the degradation induced by **1a** in both HepG_2_ and 293T cells (**Figure 4B** and **Figure S7**). Furthermore, immuno-histochemical imaging clearly showed that **1a** treatment promoted sEH colocalization with LAMP2, a lysosome marker, and co-treatment with BafA1 reversed this trend (**Figure 4C**), supporting that the sEH degradation is lysosome dependent.

**Figure 4.**
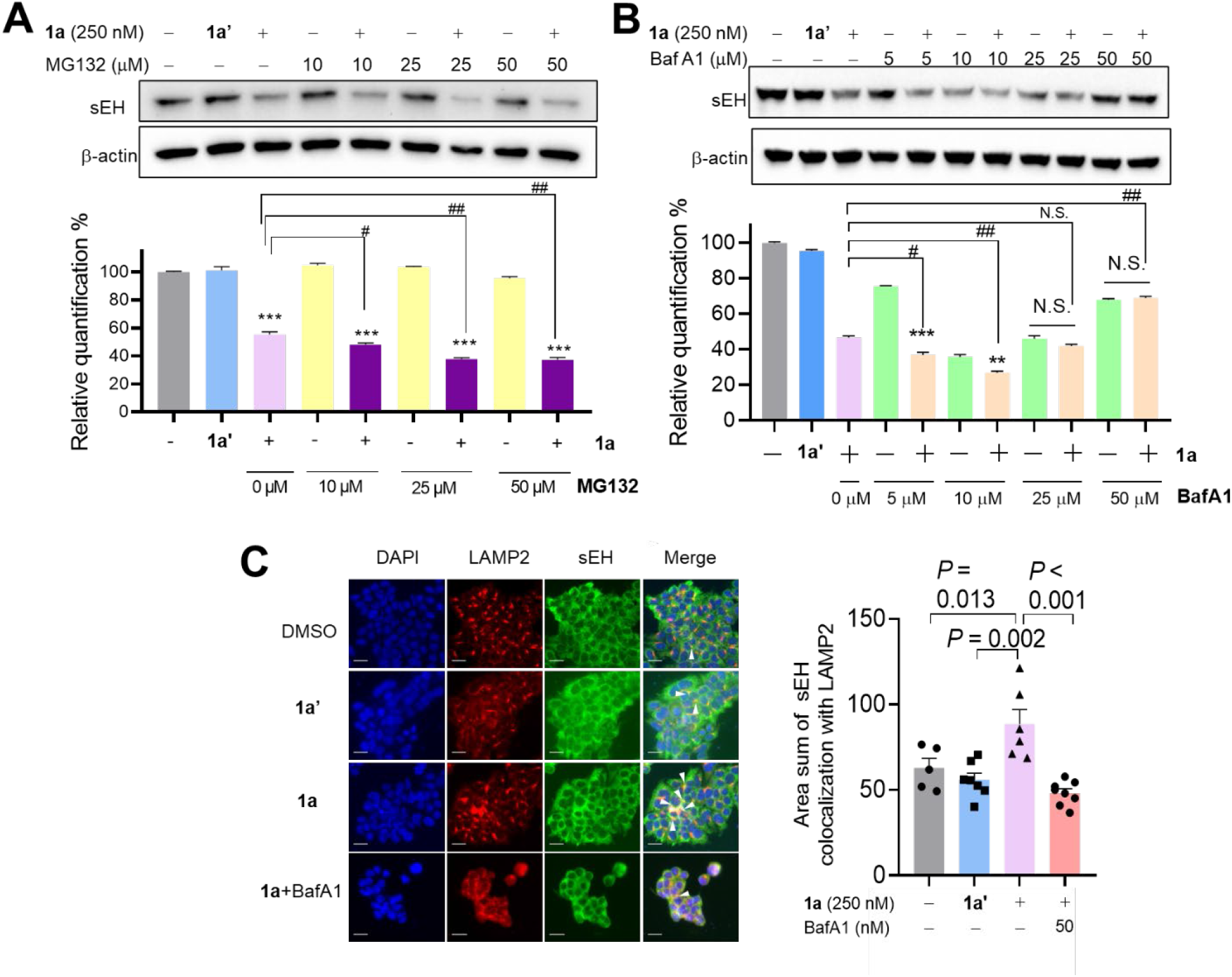
Compound 1a appears to degrade sEH through a lysosome dependent pathway. The immunoblot results of sEH levels in HepG_2_ cells treated for 24 hours with **1a** (250 nM) together with: (**A**) proteasome inhibitor MG132, or (**B**) lysosome inhibitor BafA1. Student’s t-test or Wilcoxon-Mann-Whitney test was performed depending on the normality, and the asterisks represent the significant difference between MG132 with/without **1a** or BafA1 with/without **1a** (*: *P* < 0.05, **: *P* < 0.01, ***: *P* < 0.001), the hashtags represent the significant difference between treatment vs **1a** (#: *P* < 0.05, ##: *P* < 0.01). (**C**) Immunofluorescence results of the colocalization of sEH with LAMP2. Left panel: the representative images of immunofluorescence; Right panel: the quantification results of immunofluorescence. The area of colocalization was normalized to the number of cells in the corresponding images. Scale bar, 25 mm. Mean ± SEM are shown. The difference between DMSO, **1a**’, and **1a** were analyzed using one-way ANOVA Holm Sidak test. The difference between **1a**, and **1a** with BafA1 was analyzed by Student’s t-test.

Lysosome-dependent protein degradation pathways are divided into three major classes; macroautophagy, endocytosis/microautophagy, and chaperon-mediated autophagy (Huber & Teis, 2016). We tested these previously described lysosomal pathways to determine the exact degradation mechanisms, including using ATG2A/2B ^-/-^ 293T cell or knockdown of ATG5 in 293T cell (macroautophagy), knockdown of the tumor susceptibility gene 101 (*tsg101*) (endocytosis/microautophagy), and Hsp70 family inhibitor VER155008 (microautophagy and chaperon mediated autophagy) (**Figure S8**). All these treatments failed to rescue the sEH degradation induced by **1a**, indicating that its degradation is through a previously uncharacterized lysosomal degradation pathway, or, alternatively, there is an unknown compensation mechanism for sEH degradation. Determining the exact lysosomal dependent degradation pathway is the subject of further study.

### Compound 1a has higher efficacy in reducing ER stress compared to parent sEH inhibitor 1 through inhibition of IRE1α -XBP1 signaling pathway

Previous studies demonstrated that sEH is a physiological modulator of ER stress signaling (Bettaieb, Nagata et al., 2013, Inceoglu, Bettaieb et al., 2017, Wang, Wagner et al., 2021). The efficacy of the PROTAC compounds was tested in the cellular thapsigargin (Tg)-induced ER stress assay focusing on the IRE1α -XBP1s pathway. To maximize the efficacy, we first pretreated the cells with PROTAC **1a** or **1a’** for 24 hours, then after removing the compounds, the cells were treated with Tg for 24 hours to induce ER stress. PROTAC **1a** decreased the phosphorylated IRE1α and the downstream splicing XBP1 in both HepG_2_ and 293T cells (**Figure 5A**). Consistent with immunoblot results, **1a** also rescued the cell death induced by ER stress (**Figure 5B**). In addition, compound **1a** has better efficacy on rescuing cell death when treated with Tg together for 24 hours (**Figure S9**).

**Figure 5.**
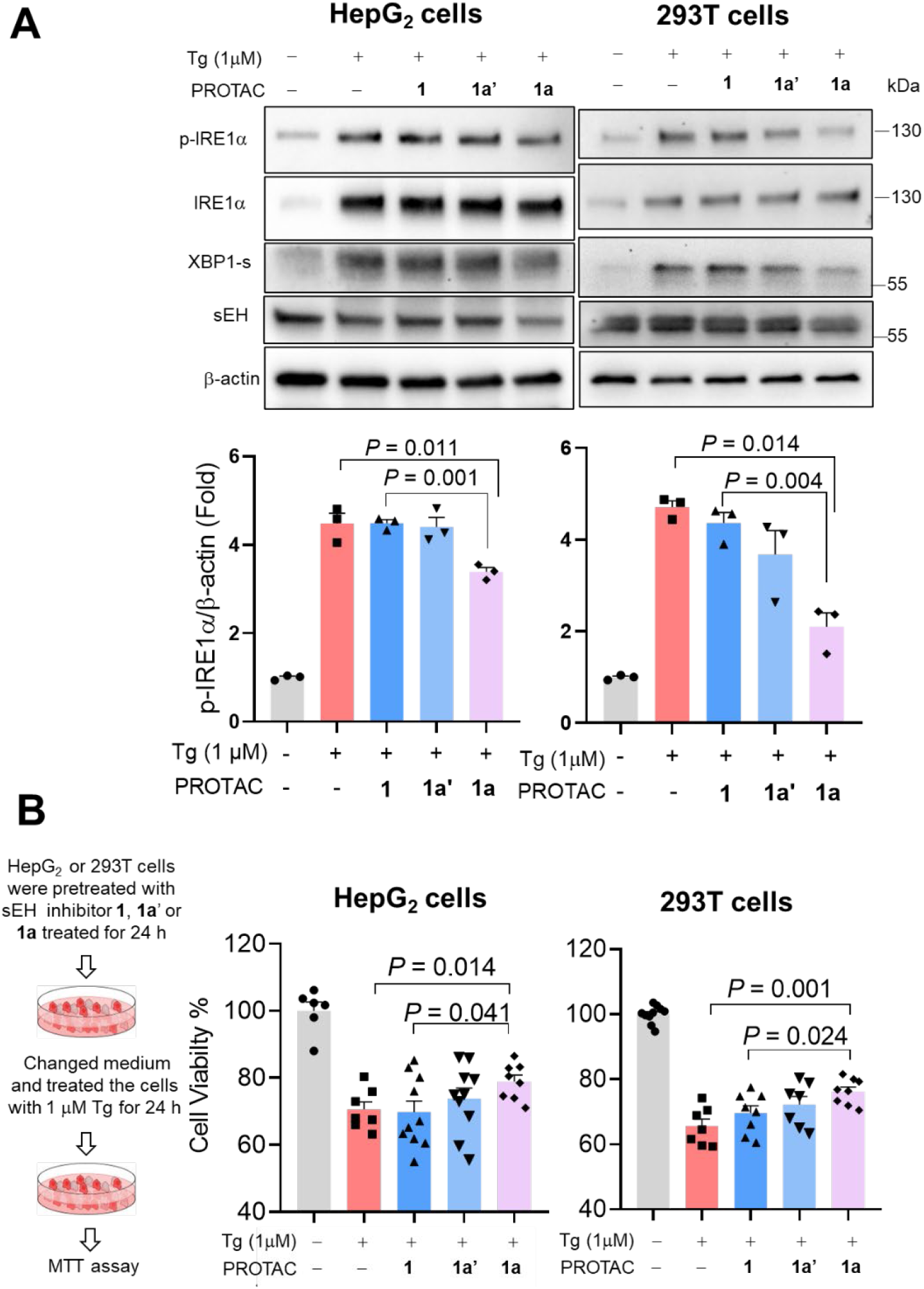
Compound 1a has higher efficacy in reducing ER stress compared to parent sEH inhibitor 1 through inhibition of IRE1α -XBP1 signaling pathway. (**A**) The HepG_2_ or 293T cells were treated with Tg, sEH inhibitor **1, 1a’**, or **1a** for 24 hours, the immunoblot results of the proteins in IRE1α-XBP1 pathway. Left panel, the representative results of immunoblot; Right panel, the quantification of immunoblot results. (**B**) The HepG_2_ or 293T cells were pre-treated with sEH inhibitor **1, 1a’**, or **1a** (250 nM) for 24 hours, then changed the medium containing Tg without PROTACs and incubated the cells for another 24 hours. The cell death was measured using MTT assay. Left panel, the procedure scheme of the treatment; Right panel, MTT assay results. n = 6-10. Tg, thapsigargin. Mean ± SEM are shown. The difference between two groups was analyzed by Student’s t-test or Wilcoxon-Mann-Whitney depending on the normality.

## Discussion

We here report the development of the first-in-class small molecule degraders of sEH and their application to study the biology of sEH. For the development of the PROTAC molecules, we employed two scaffolds of sEH inhibitors, AUDA and *t*-TUCB (**Table 1**). AUDA is one of the first generation sEH inhibitors with high potency against most mammalian sEH homologs while *t*-TUCB is a newer inhibitor with improved physicochemical properties and metabolic stability (Hwang et al., 2007, Morisseau, Goodrow et al., 2002). In both cases, previous SAR and x-ray complex structures showed that carboxylic acid is not required for potency. Therefore, we connected the carboxylic acid to the recruiters of widely used E3 ligases (Von Hippel-Lindau (VHL) or cereblon (CRBN)) with various linker lengths and types (alkyl chain or PEG) to optimize the degradation potency.

For the prioritization of the synthesized molecules, four complementary assays were used to detect the sEH levels. Globally, most of the assays showed similar trend of sEH degradation. For example, we observed a high correlation (r = 0.82) between the EH-hydrolase activity and the band intensity in immunoblotting. Interestingly, there are a few discrepancies among the data from these four methods. For example, compound **1e** showed high degradation effects as monitored by the immunoblotting and sEH hydrolase activity, while only the limited effects were observed in the phosphatase activity and ELISA. Determining the exact mechanism of this interesting observation is the subject of further study. For future screening and optimization of sEH PROTACs, the ELISA assay will be the method of choice given the throughput, robustness, and quantitative aspects. Nevertheless, in all assays, compound **1a** showed high degradation efficacy and this compound was selected for further biological characterization.

The PROTAC decreased the enzyme activity of both hydrolase and phosphatase activities (**Figure 1B**). The phosphatase domain has been shown to metabolize the lysophosphatidic acids (Morisseau et al., 2012), regulate the subcellular localization of sEH (Kramer & Proschak, 2017), and negatively regulate simvastatin-activated eNOS by impeding the Akt–AMPK–eNOS signaling cascade (Hou, Liao et al., 2015). The development of potent small-molecule phosphatase inhibitors has been challenging (Kramer, Woltersdorf et al., 2019). As an alternative approach, the molecules developed here could be useful chemical probes to study the role of sEH phosphatase activity.

One of our central findings is that sEH PROTAC selectively instigates the degradation of cytosolic sEH while peroxisomal sEH is resistant to degradation. It is reported that peroxisomal translocation of soluble epoxide hydrolase protects against ischemic stroke injury (Nelson et al., 2015), and sEH dimerization status is a key regulator of its peroxisomal localization (Nelson, Das et al., 2016). The sEH PROTAC compounds will be useful chemical probes to understand the functional difference of peroxisomal vs cytosolic sEH. Furthermore, our study showcases the proof of concept of selective modulation of the protein function in a subcellular-component specific manner using PROTACs. Cytosol selective degradation is likely due to the inaccessibility of the E3-ligase complex and/or proteasome/lysosome to the inside of the peroxisome. So far, PROTACs have been developed to target cytosolic proteins, nuclear proteins, endosomal membrane proteins, single- and multi-pass transmembrane proteins (Bai et al., 2019, Bond et al., 2020, Neklesa et al., 2019), while PROTACs have not been reported to target mitochondrial proteins, ER-localized proteins, and Golgi-localized proteins. All these proteins that are not degradable by PROTACs are within a membrane-surrounded subcellular compartment. Developing small molecule degraders of these proteins, including sEH in peroxisome, is a challenge for future research.

The sEH PROTAC compounds are thalidomide- or VHL ligand-based PRTOACs that induce protein degradation through the ubiquitin-proteasome system (UPS) (Bondeson, Smith et al., 2018, Smith, Wang et al., 2019). However, interestingly in our case, co-treatment of proteasome inhibitors with compound **1a** did not rescue sEH (**Figures 4A**). Surprisingly, lysosomal inhibitor BafA1 rescued sEH, suggesting that sEH is degraded through the lysosomal pathway (**Figures 4C** and **S6**). Further research is needed to determine the exact mechanisms of sEH degradation induced by compound **1a** as well as endogenous degradation pathway. Although our mechanistic data clearly suggest that the binding of our molecules on sEH and E3 ligase is essential for degradation (**Figures S2** and **S3**), we cannot exclude the possibility that the degradation is caused by secondary effects or off-target effect at this point. The sEH PROTAC compound described herein could be the first PROTAC that induces the degradation of a soluble target protein through the lysosome. A recent study showed that PROTACs targeting a membrane protein EGFR also degrade through a lysosome-dependent manner (Qu, Liu et al., 2021), which indicates some of the PROTACs may induce proteasome independent and lysosome dependent degradation. Based on this, it is strongly recommended to test the degradation mechanism of PROTACs using proteasome inhibitors. Endogenous ubiquitination of proteins sometimes leads to lysosomal degradation, while the exact determinant of the specific degradation mechanism is still unclear. For example, it is known that some long-lived proteins are endogenously degraded through the lysosome. Consistent with previous proteomic analysis, the cellular turnover of sEH appeared to be slow (**Figure 3**) (Mathieson, Franken et al., 2018). At this point, it is unclear whether the lysosome-dependent degradation is the only degradation mechanism of sEH. Our findings, together with the study on PROTAC-induced lysosome-dependent EGFR degradation (Qu et al., 2021), suggest that the PROTACs can also induce lysosome-dependent degradation of target proteins and caution the current assumption of the mechanism of action of PROTACs.

Finally, we demonstrated that the sEH PROTAC compound has enhanced potency compared to the parent inhibitor using a cellular ER stress assay. This is likely due to the PROTACs-induced sustained degradation of sEH during ER stress. This could be advantageous for clinical applications through reducing undesirable side effects with prolonged efficacy. Determining the *in vivo* efficacy of the developed PROTACs is the subject of further study.

In summary, the sEH PROTAC compounds reported herein selectively degrade cytosolic sEH and not peroxisomal sEH, providing a useful tool to study the function of sEH in a spatiotemporal manner. Furthermore, the data indicate that our PROTAC compound induces lysosome-dependent, proteasome-independent degradation of the targeted protein, shedding light on the lysosomal-dependent degradation pathway induced by PROTACs and questioning the current mechanistic assumption of PROTACs.

## Supporting information

Original western data

## Acknowledgments

We thank Dr. Ana Cristina G Grodzki for assistance with immunofluorescence, and grant P50HD103526 from IDDRC-MIND Institute, UC Davis for the use of the imaging facility from the Biological and Molecular Analysis Core (BMAC). We would like to thank Dr. Ian Wilson and Dr. K. Barry Sharpless for support and encouragement on this project and access to the instrument. We also thank Dr. Komatsu Masaaki for kindly providing the ATG 2A/2B^-/-^ cells, Dr. Sung Hee Hwang for kindly providing the sEH inhibitors *t*-TUCB and AUDA, and Dr. Natalia Orendain for her help on statistical data analysis. We would also like to thank Dr. Takanori Otomo for helpful discussion and help on obtaining cells and project management. This study was supported, in part, by a grant from the National Institute of Environmental Health Sciences (NIEHS) Grant R35ES030443 (B.D.H.), NIEHS Superfund Research Program P42 ES004699 (B.D.H.), and National Institute of General Medical Sciences of the National Institutes of Health under Award Number R35GM136286 (D.W.W.), K99GM138758 & R00GM138758 (S.K.). The Sidoli lab gratefully acknowledges for financial support AFAR (Sagol Network GerOmics award), Deerfield (Xseed award), Relay Therapeutics, Merck, the NIH Office of the Director (1-S10-OD030286-01), the Einstein-Mount Sinai Diabetes Research Center, and the Einstein Cancer Center (P30-CA013330-47).

## Author contributions

Conceptualization: Y.W., C.M., S.K.; Synthesis: S.K., A.T.; Data acquisition and analysis: Y.W., C.M., D.W., S.S., J.Y., D.L., S.K.; Visualization: Y.W. and S.K.; Project administration: C.M., S. K.; Writing—original draft preparation: Y.W., S.K. Supervision: D.W.W., B.D.H.. All authors have read, reviewed, edited, and agreed to the published version of the manuscript.

## References

Aguilan JT, Kulej K, Sidoli S (2020) Guide for protein fold change and p-value calculation for non-experts in proteomics. Mol Omics 16: 573–582

Bai L, Zhou H, Xu R, Zhao Y, Chinnaswamy K, McEachern D, Chen J, Yang CY, Liu Z, Wang M, Liu L, Jiang H, Wen B, Kumar P, Meagher JL, Sun D, Stuckey JA, Wang S (2019) A Potent and Selective Small-Molecule Degrader of STAT3 Achieves Complete Tumor Regression In Vivo. Cancer Cell 36: 498–511 e17

Bettaieb A, Nagata N, AbouBechara D, Chahed S, Morisseau C, Hammock BD, Haj FG (2013) Soluble epoxide hydrolase deficiency or inhibition attenuates diet-induced endoplasmic reticulum stress in liver and adipose tissue. J Biol Chem 288: 14189–14199

Bond MJ, Chu L, Nalawansha DA, Li K, Crews CM (2020) Targeted Degradation of Oncogenic KRAS(G12C) by VHL-Recruiting PROTACs. ACS Cent Sci 6: 1367–1375

Bondeson DP, Smith BE, Burslem GM, Buhimschi AD, Hines J, Jaime-Figueroa S, Wang J, Hamman BD, Ishchenko A, Crews CM (2018) Lessons in PROTAC Design from Selective Degradation with a Promiscuous Warhead. Cell Chem Biol 25: 78–87 e5

Borhan B, Mebrahtu T, Nazarian S, Kurth MJ, Hammock BD (1995) Improved radiolabeled substrates for soluble epoxide hydrolase. Anal Biochem 231: 188–200

Enayetallah AE, French RA, Barber M, Grant DF (2006) Cell-specific subcellular localization of soluble epoxide hydrolase in human tissues. J Histochem Cytochem 54: 329–35

Fishbein A, Wang W, Yang H, Yang J, Hallisey VM, Deng J, Verheul SML, Hwang SH, Gartung A, Wang Y, Bielenberg DR, Huang S, Kieran MW, Hammock BD, Panigrahy D (2020) Resolution of eicosanoid/cytokine storm prevents carcinogen and inflammation-initiated hepatocellular cancer progression. Proc Natl Acad Sci U S A 117: 21576–21587

Gao H, Sun X, Rao Y (2020) PROTAC Technology: Opportunities and Challenges. ACS Med Chem Lett 11: 237–240

Garnar-Wortzel L, Bishop TR, Kitamura S, Milosevich N, Asiaban JN, Zhang X, Zheng Q, Chen E, Ramos AR, Ackerman CJ, Hampton EN, Chatterjee AK, Young TS, Hull MV, Sharpless KB, Cravatt BF, Wolan DW, Erb MA (2021) Chemical Inhibition of ENL/AF9 YEATS Domains in Acute Leukemia. ACS Cent Sci 7: 815–830

Gill S, Grant DF, Beetham JK, Chang C, Hammock BD (1994) Peroxisomal proliferation and subcellular localization of soluble epoxide hydrolase: Role of peroxisomal targeting sequences. Peroxisome Proliferators: Unique Inducers of Drug-Metabolizing Enzymes: 113–121

Hashimoto K (2019) Role of Soluble Epoxide Hydrolase in Metabolism of PUFAs in Psychiatric and Neurological Disorders. Front Pharmacol 10: 36

Hou HH, Liao YJ, Hsiao SH, Shyue SK, Lee TS (2015) Role of phosphatase activity of soluble epoxide hydrolase in regulating simvastatin-activated endothelial nitric oxide synthase. Sci Rep 5: 13524

Huber LA, Teis D (2016) Lysosomal signaling in control of degradation pathways. Curr Opin Cell Biol 39: 8–14

Hwang SH, Tsai HJ, Liu JY, Morisseau C, Hammock BD (2007) Orally bioavailable potent soluble epoxide hydrolase inhibitors. J Med Chem 50: 3825–40

Imig JD, Hammock BD (2009) Soluble epoxide hydrolase as a therapeutic target for cardiovascular diseases. Nat Rev Drug Discov 8: 794–805

Imig JD, Zhao X, Capdevila JH, Morisseau C, Hammock BD (2002) Soluble epoxide hydrolase inhibition lowers arterial blood pressure in angiotensin II hypertension. Hypertension 39: 690–4

Inceoglu B, Bettaieb A, Haj FG, Gomes AV, Hammock BD (2017) Modulation of mitochondrial dysfunction and endoplasmic reticulum stress are key mechanisms for the wide-ranging actions of epoxy fatty acids and soluble epoxide hydrolase inhibitors. Prostaglandins Other Lipid Mediat 133: 68–78

Ji CH, Kwon YT (2017) Crosstalk and Interplay between the Ubiquitin-Proteasome System and Autophagy. Mol Cells 40: 441–449

Jones PD, Wolf NM, Morisseau C, Whetstone P, Hock B, Hammock BD (2005) Fluorescent substrates for soluble epoxide hydrolase and application to inhibition studies. Anal Biochem 343: 66–75

Kitamura S, Morisseau C, Harris TR, Inceoglu B, Hammock BD (2017) Occurrence of urea-based soluble epoxide hydrolase inhibitors from the plants in the order Brassicales. PLoS One 12: e0176571

Kitamura S, Morisseau C, Inceoglu B, Kamita SG, De Nicola GR, Nyegue M, Hammock BD (2015) Potent natural soluble epoxide hydrolase inhibitors from Pentadiplandra brazzeana baillon: synthesis, quantification, and measurement of biological activities in vitro and in vivo. PLoS One 10: e0117438

Kramer J, Proschak E (2017) Phosphatase activity of soluble epoxide hydrolase. Prostaglandins Other Lipid Mediat 133: 88–92

Kramer JS, Woltersdorf S, Duflot T, Hiesinger K, Lillich FF, Knoll F, Wittmann SK, Klingler FM, Brunst S, Chaikuad A, Morisseau C, Hammock BD, Buccellati C, Sala A, Rovati GE, Leuillier M, Fraineau S, Rondeaux J, Hernandez-Olmos V, Heering J et al. (2019) Discovery of the First in Vivo Active Inhibitors of the Soluble Epoxide Hydrolase Phosphatase Domain. J Med Chem 62: 8443–8460

Lai AC, Crews CM (2017) Induced protein degradation: an emerging drug discovery paradigm. Nat Rev Drug Discov 16: 101–114

Li D, Cui Y, Morisseau C, Gee SJ, Bever CS, Liu X, Wu J, Hammock BD, Ying Y (2017) Nanobody Based Immunoassay for Human Soluble Epoxide Hydrolase Detection Using Polymeric Horseradish Peroxidase (PolyHRP) for Signal Enhancement: The Rediscovery of PolyHRP? Anal Chem 89: 6248–6256

Liu J, Chen H, Ma L, He Z, Wang D, Liu Y, Lin Q, Zhang T, Gray N, Kaniskan HU, Jin J, Wei W (2020) Light-induced control of protein destruction by opto-PROTAC. Sci Adv 6: eaay5154

Mathieson T, Franken H, Kosinski J, Kurzawa N, Zinn N, Sweetman G, Poeckel D, Ratnu VS, Schramm M, Becher I, Steidel M, Noh KM, Bergamini G, Beck M, Bantscheff M, Savitski MM (2018) Systematic analysis of protein turnover in primary cells. Nat Commun 9: 689

Moody DE, Loury DN, Hammock BD (1985) Epoxide metabolism in the liver of mice treated with clofibrate (ethyl-alpha-(p-chlorophenoxyisobutyrate)), a peroxisome proliferator. Toxicol Appl Pharmacol 78: 351–62

Morisseau C, Bernay M, Escaich A, Sanborn JR, Lango J, Hammock BD (2011) Development of fluorescent substrates for microsomal epoxide hydrolase and application to inhibition studies. Anal Biochem 414: 154–62

Morisseau C, Goodrow MH, Dowdy D, Zheng J, Greene JF, Sanborn JR, Hammock BD (1999) Potent urea and carbamate inhibitors of soluble epoxide hydrolases. Proc Natl Acad Sci U S A 96: 8849–54

Morisseau C, Goodrow MH, Newman JW, Wheelock CE, Dowdy DL, Hammock BD (2002) Structural refinement of inhibitors of urea-based soluble epoxide hydrolases. Biochem Pharmacol 63: 1599–608

Morisseau C, Hammock BD (2013) Impact of soluble epoxide hydrolase and epoxyeicosanoids on human health. Annu Rev Pharmacol Toxicol 53: 37–58

Morisseau C, Schebb NH, Dong H, Ulu A, Aronov PA, Hammock BD (2012) Role of soluble epoxide hydrolase phosphatase activity in the metabolism of lysophosphatidic acids. Biochem Biophys Res Commun 419: 796–800

Neklesa T, Snyder LB, Willard RR, Vitale N, Pizzano J, Gordon DA, Bookbinder M, Macaluso J, Dong H, Ferraro C, Wang G, Wang J, Crews CM, Houston J, Crew AP, Taylor I (2019) ARV-110: An oral androgen receptor PROTAC degrader for prostate cancer. Journal of Clinical Oncology 37: 259–259

Nelson JW, Das AJ, Barnes AP, Alkayed NJ (2016) Disrupting Dimerization Translocates Soluble Epoxide Hydrolase to Peroxisomes. PLoS One 11: e0152742

Nelson JW, Zhang W, Alkayed NJ, Koerner IP (2015) Peroxisomal translocation of soluble epoxide hydrolase protects against ischemic stroke injury. J Cereb Blood Flow Metab 35: 1416–20

Newman JW, Morisseau C, Harris TR, Hammock BD (2003) The soluble epoxide hydrolase encoded by EPXH2 is a bifunctional enzyme with novel lipid phosphate phosphatase activity. Proc Natl Acad Sci U S A 100: 1558–63

Pettersson M, Crews CM (2019) PROteolysis TArgeting Chimeras (PROTACs) - Past, present and future. Drug Discov Today Technol 31: 15–27

Qu X, Liu H, Song X, Sun N, Zhong H, Qiu X, Yang X, Jiang B (2021) Effective degradation of EGFR(L858R+T790M) mutant proteins by CRBN-based PROTACs through both proteosome and autophagy/lysosome degradation systems. Eur J Med Chem 218: 113328

Sakamoto KM, Kim KB, Kumagai A, Mercurio F, Crews CM, Deshaies RJ (2001) Protacs: chimeric molecules that target proteins to the Skp1-Cullin-F box complex for ubiquitination and degradation. Proc Natl Acad Sci U S A 98: 8554–9

Salami J, Alabi S, Willard RR, Vitale NJ, Wang J, Dong H, Jin M, McDonnell DP, Crew AP, Neklesa TK, Crews CM (2018) Androgen receptor degradation by the proteolysis-targeting chimera ARCC-4 outperforms enzalutamide in cellular models of prostate cancer drug resistance. Commun Biol 1: 100

Schneider-Poetsch T, Ju J, Eyler DE, Dang Y, Bhat S, Merrick WC, Green R, Shen B, Liu JO (2010) Inhibition of eukaryotic translation elongation by cycloheximide and lactimidomycin. Nat Chem Biol 6: 209–217

Smith BE, Wang SL, Jaime-Figueroa S, Harbin A, Wang J, Hamman BD, Crews CM (2019) Differential PROTAC substrate specificity dictated by orientation of recruited E3 ligase. Nat Commun 10: 131

Sun X, Gao H, Yang Y, He M, Wu Y, Song Y, Tong Y, Rao Y (2019) PROTACs: great opportunities for academia and industry. Signal Transduct Target Ther 4: 64

Wagner KM, McReynolds CB, Schmidt WK, Hammock BD (2017) Soluble epoxide hydrolase as a therapeutic target for pain, inflammatory and neurodegenerative diseases. Pharmacol Ther 180: 62–76

Wang W, Yang J, Zhang J, Wang Y, Hwang SH, Qi W, Wan D, Kim D, Sun J, Sanidad KZ, Yang H, Park Y, Liu JY, Zhao X, Zheng X, Liu Z, Hammock BD, Zhang G (2018) Lipidomic profiling reveals soluble epoxide hydrolase as a therapeutic target of obesity-induced colonic inflammation. Proc Natl Acad Sci U S A 115: 5283–5288

Wang Y, Wagner KM, Morisseau C, Hammock BD (2021) Inhibition of the Soluble Epoxide Hydrolase as an Analgesic Strategy: A Review of Preclinical Evidence. J Pain Res 14: 61–72

Zou Y, Ma D, Wang Y (2019) The PROTAC technology in drug development. Cell Biochem Funct 37: 21–30

