## Supporting Information for "PROTAC-mediated selective degradation of cytosolic soluble epoxide hydrolase enhances ER-stress reduction"

### Table of contents

**Tables**

**Table S1.** The molecular weight and inhibitory potency of PROTAC compounds against human and mouse sEH.
**Table S2.** The *P* values in Figure 1B.

**Figures**

**Fig. S1.** Dose response of sEH PROTAC compounds **1e** and **2a** in $HepG_2$ cells.
**Fig. S2.** sEH binding of PROTAC **1a** is required for the degradation effect.
**Fig. S3.** The effect of MLN4924 on sEH degradation induced by PROTAC **1a** in $HepG_2$ cells.
**Fig. S4.** The degradation effect of **1a** (250 nM) in $HepG_2$ cells for 24 or 48 hours.
**Fig. S5.** Endogenous degradation of sEH is not inhibited by the proteasome inhibitor BTZ.
**Fig. S6.** The effect of BTZ on sEH degradation induced by **1a**, **1e**, or **2a** in $HepG_2$ cells.
**Fig. S7.** The effect of BafA1 on sEH degradation induced by PROTAC **1a** in 293T cells.
**Fig. S8.** The degradation mechanism of sEH PROTAC **1a**.
**Fig. S9.** The effect of compound **1a** on Tg-induced cell viability in $HepG_2$ cells.

**Synthetic methods and compound characterization**

**Table S1.** The molecular weight and inhibitory potency of PROTAC compounds against human and mouse sEH

| Compound number | Molecular Weight | Human sEH $IC_{90}$ (nM) | Mouse sEH $IC_{50}$ (nM) | Mouse sEH $IC_{90}$ (nM) | Mouse sEH $IC_{50}$ phosphatase (nM) |
|---|---|---|---|---|---|
| **1** | 438.4 | 2.7 | 0.7 | 14.6 | > 10,000 |
| **1a'** | 985.0 | 8.8 | 1.0 | 20.6 | |
| **1a** | 971.0 | 12.2 | 5.5 | 66.7 | |
| **1b** | 822.8 | 4.8 | 2.7 | 38.2 | |
| **1c** | 882.9 | 4.4 | 1.2 | 13.3 | |
| **1d** | 850.8 | 6.1 | 0.9 | 21.7 | |
| **1e** | 978.1 | 6.7 | 2.1 | 60.2 | |
| **1f** | 922.0 | 19.1 | 11.6 | 180.2 | |
| **1g** | 996.1 | 7.3 | 3.8 | 47.3 | |
| **2** | 392.6 | 8.1 | 0.5 | 7.8 | |
| **2a'** | 939.2 | 5.1 | 0.7 | 7.4 | |
| **2a** | 925.1 | 12.9 | 1.8 | 10.8 | |
| **2b** | 777.0 | 4.5 | 1.3 | 7.6 | |
| **2c** | 837.0 | 6.8 | 2.2 | 19.7 | |
| **2d** | 805.0 | 6.8 | 2.2 | 19.7 | |
| **2e** | 932.3 | 5.5 | 0.9 | 4.4 | |
| **2f** | 876.2 | 5.8 | 0.9 | 4.9 | |
| **2g** | 950.3 | 6.3 | 2.0 | 10.6 | |

**Table S2.** The *P* values in Figure 1B

| Compound | Comparison with | *P* value | | | t, ANOVA |
|---|---|---|---|---|---|
| | | Hydrolase activity | Phosphatase activity | ELISA | |
| **1** | DMSO | N.S. | N.S. | N.S. | t-test |
| **1a'** | DMSO | 0.010 | 0.007 | N.S. | t-test |
| **1a** | DMSO | <0.001 | <0.001 | 0.012 | ANOVA |
| | **1** | 0.010 | <0.001 | N.S. | ANOVA |
| | **1a'** | <0.001 | <0.001 | N.S. | ANOVA |
| **1c** | DMSO | N.S. | N.S. | N.S. | t-test |
| | **1** | N.S. | N.S. | N.S. | t-test |
| | **1a'** | N.S. | N.S. | N.S. | t-test |
| **1e** | DMSO | <0.001 | 0.005 | N.S. | ANOVA |
| | **1** | 0.006 | 0.006 | N.S. | ANOVA |
| | **1a'** | N.S. | N.S. | N.S. | ANOVA |
| **2** | DMSO | 0.017 | N.S. | N.S. | t-test |
| **2a'** | DMSO | 0.026 | N.S. | N.S. | t-test |
| **2a** | DMSO | 0.005 | 0.002 | <0.001 | ANOVA |
| | **2** | N.S. | 0.007 | <0.001 | ANOVA |
| | **2a'** | N.S. | <0.001 | <0.001 | ANOVA |
| **2c** | DMSO | 0.004 | 0.017 | <0.001 | ANOVA |
| | **2** | N.S. | 0.009 | <0.001 | ANOVA |
| | **2a'** | N.S. | 0.011 | <0.001 | ANOVA |
| **2d** | DMSO | 0.003 | 0.002 | 0.004 | ANOVA |
| | **2** | N.S. | 0.009 | N.S. | ANOVA |
| | **2a'** | N.S. | <0.001 | 0.008 | ANOVA |
| **2e** | DMSO | 0.005 | 0.002 | <0.001 | ANOVA |
| | **2** | N.S. | 0.009 | <0.001 | ANOVA |
| | **2a'** | N.S. | <0.001 | <0.001 | ANOVA |

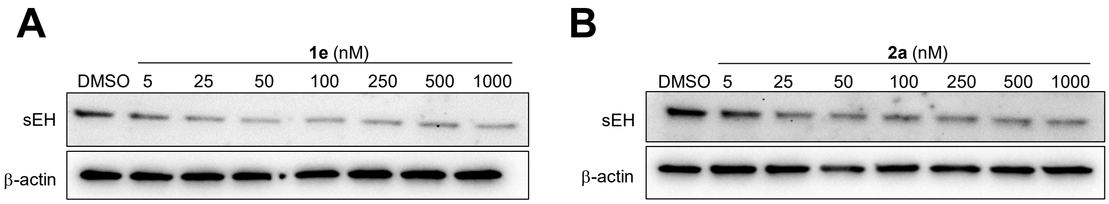


**Figure S1. Dose response of sEH PROTAC compounds 1e and 2a in HepG₂ cells.** The concentration response of **(A) 1e**, **(B) 2a**. HepG₂ cells were treated with indicated concentration of **1e,** or **2a** for 24 hours and sEH level was measured by immunoblot.

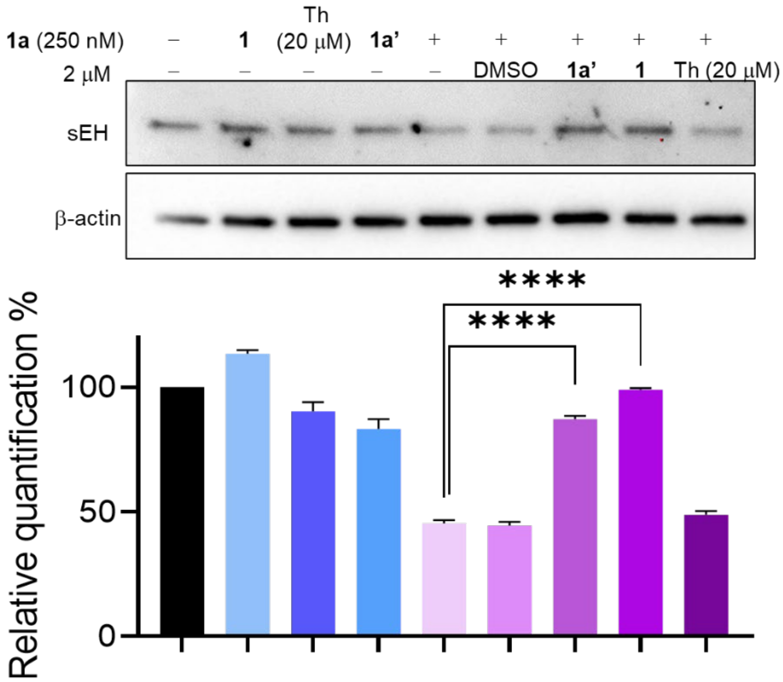


**Figure S2. sEH binding of PROTAC 1a is required for the degradation effect.** The HepG₂ cells were treated as indicated in the figure and the cells were collected for immunoblot analysis. Th; thalidomide. Student's t-test was performed, and the asterisks represent the significant difference between treatment vs **1a** (****: $P < 0.001$).

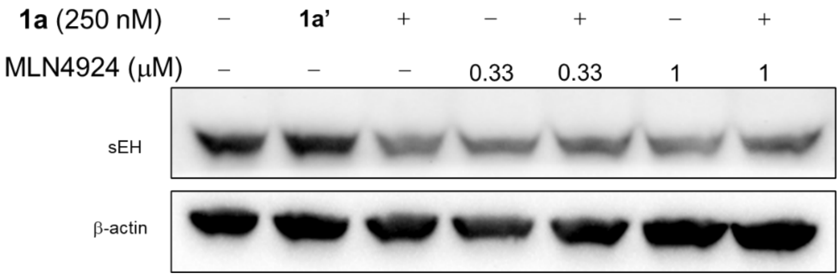


**Figure S3. The effect of MLN4924 on sEH degradation induced by PROTAC 1a in HepG$_2$ cells.** The HepG$_2$ cells were treated with DMSO, PROTAC **1a**, and indicated concentration of MLN4924 for 24 hours, then the cells were collected for immunoblot analysis.

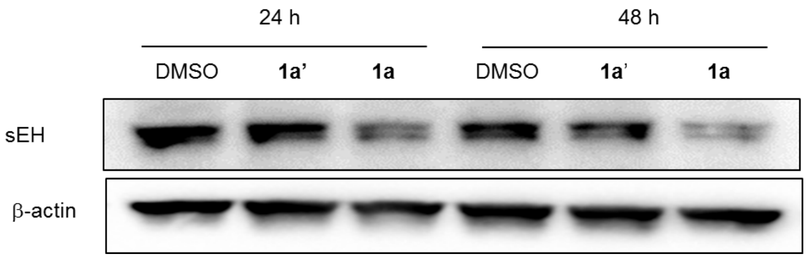


**Figure S4. The degradation effect of 1a (250 nM) in HepG$_2$ cells for 24 or 48 hours.**

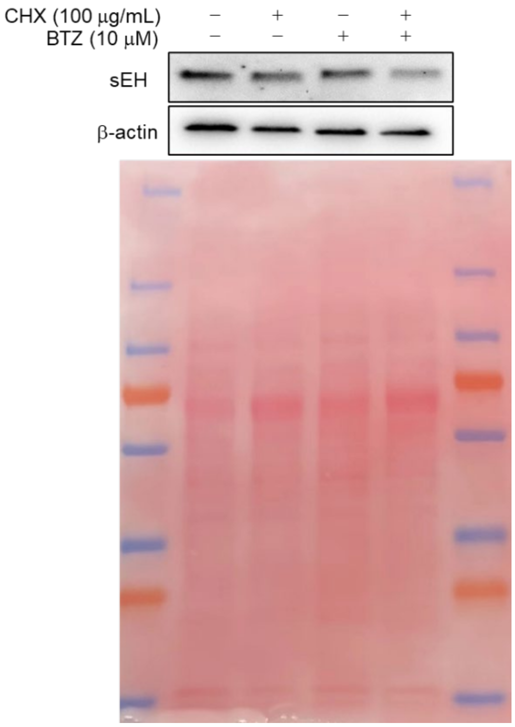


**Figure S5. Endogenous degradation of sEH is not inhibited by the proteasome inhibitor BTZ.** $HepG_2$ cells were treated with CHX with/without BTZ for 48 hours and sEH was detected using immunoblot. Top panel, the representative results of immunoblot; Bottom panel, the quantification of immunoblot results. The whole membrane staining had been shown here as a loading control since β-actin level decreased after 48 h treatment of CHX.

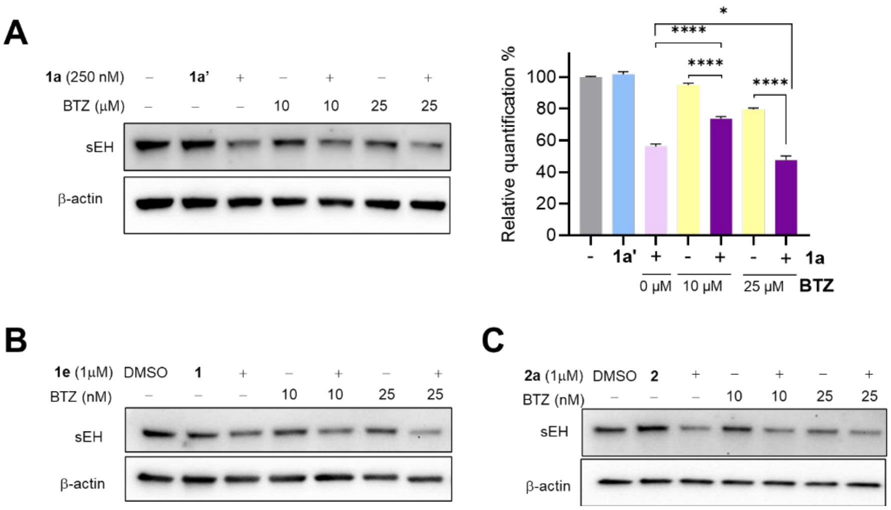


**Figure S6. The effect of BTZ on sEH degradation induced by 1a, 1e, or 2a in $HepG_2$ cells.** The $HepG_2$ cells were treated with DMSO, PROTAC **1a** (**A**), PROTAC **1e** (**B**), PROTAC **2a** (**C**), and indicated concentration of BTZ for 24 hours, then the cells were collected for immunoblot analysis.

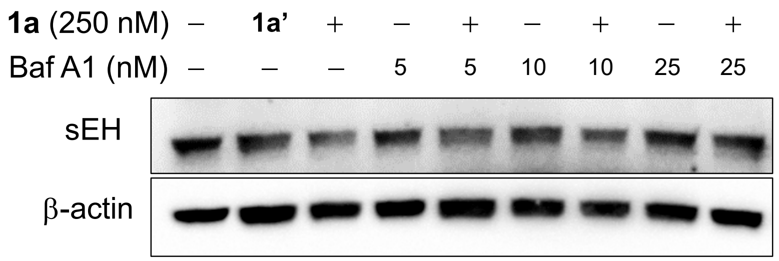


**Figure S7. The effect of BafA1 on sEH degradation induced by PROTAC 1a in 293T cells.** The 293T cells were treated with DMSO, PROTAC **1a'**, PROTAC **1a**, and indicated concentration of BafA1 for 24 hours, then the cells were collected for immunoblot analysis.

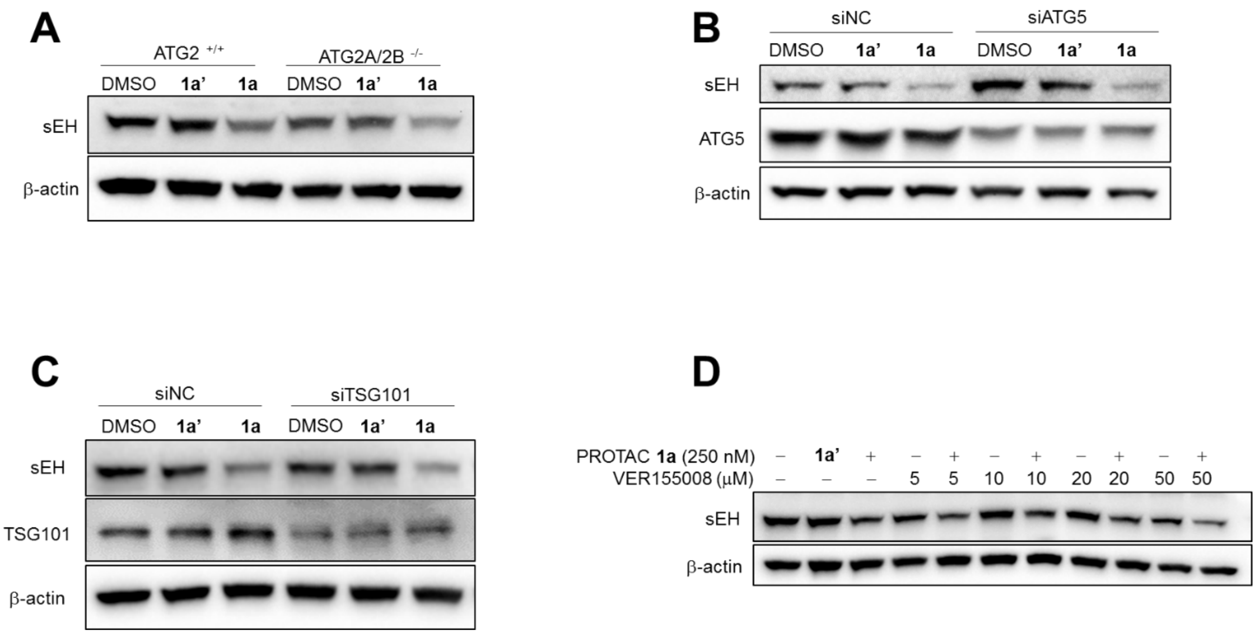


**Figure S8. The degradation mechanism of sEH PROTAC 1a.** (**A**) The effect of **1a** in wild type 293T and ATG 2A/2B $^{-/-}$ cells. (**B**) 293T cells were transfected with negative control or siATG5 for 24 hours, then the cells were treated with DMSO, **1a'** or **1a** for 24 hours, then the cells were collected for immunoblot analysis. (**C**) 293T cells were transfected with negative control or siTsg101 for 24 hours, then the cells were treated with DMSO, **1a'** or **1a** for 24 hours, then the cells were collected for immunoblot analysis. (**D**) $HepG_2$ cells were treated with DMSO, **1a'**, or **1a**, and the indicated concentration of Hsp70 family inhibitor VER155008 for 24 hours, then the cells were collected for immunoblot analysis.

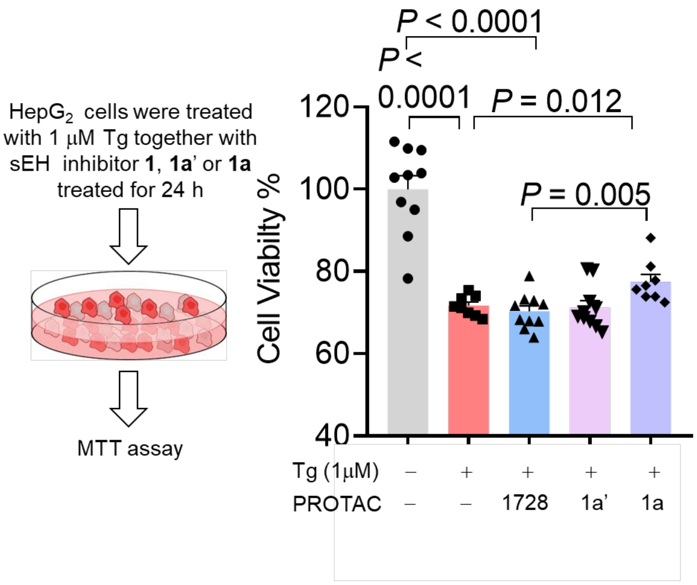


**Figure S9. The effect of compound 1a on Tg-induced cell viability in HepG$_2$ cells.** The HepG$_2$ were treated with 1 μM Tg, sEH inhibitor **1**, **1a'**, or **1a** (250 nM) for 24 hours, then the cell death was measured using MTT assay. Left panel, the procedure scheme of the treatment; Right panel, MTT assay results. n = 8-10. Tg, thapsigargin. Mean ± SEM are shown. The difference between two groups was analyzed by Student's t-test.

**Synthetic methods and compound characterization**
**General**
All reagents and solvents were purchased from commercial suppliers and were used without further purification. $^1$H and $^{13}$C NMR spectra were collected using a Bruker 600, 500, or 400 MHz spectrometer with chemical shifts reported relative to residual deuterated solvent peaks or a tetramethylsilane internal standard. $CFCl_3$ was used as an internal standard for $^{19}$F-NMR. Accurate masses were measured using Agilent G6125 mass spectrometer. Reactions were monitored on TLC plates (silica gel 60, F254 coating, EMD Millipore, 1057150001), and spots were either monitored under UV light (254 mm) or stained with phosphomolybdic acid. The same TLC system was used to test purity, and all final products showed a single spot on TLC with both $KMnO_4$ and UV absorbance. The purity of the compounds that were tested in the assay was >95% based on $^1$H NMR and reverse phase HPLC-UV on monitoring absorption at 230 nm. The HPLC gradient method consisted of an aqueous phase (Milli-Q water with 0.05% trifluoroacetic acid) and an organic phase (acetonitrile with 0.05% trifluoroacetic acid) with a 0.55 mL/min flow. The first step consisted of 90% aqueous and 10% organic phases followed by a 5-min gradient to 90% organic phase. A subsequent 1-min step of 90% organic phase was followed by a 0.5-min gradient to 90% aqueous and 10% organic phases.

**Synthesis procedure for PROTAC molecules**
Each sEH binder (*t*-TUCB or AUDA, 1.2 eq) and HATU (1.5 eq.) were added to a solution of each recruiter molecule (thalidomide or VHL ligand tethered with $NH_2$-linker, 1 eq. (Tocris Bioscience)) in DMF (16 mM) and DIPEA (10 eq.). The reaction mixture was stirred at room temperature for 1 hour, then filtered through a PTFE membrane and purified via preparative HPLC (C18, 19 x 150 mm) using a $H_2O$-MeCN gradient (99:1 to 20:80, v/v, 0.1% TFA). A fraction containing each sEH-PROTAC molecule was lyophilized to give a white solid.
For a recruiter molecule tethered with Boc-NH-linker, Boc group was deprotected using conc. HCl in 1,4-dioxane at room temperature for 10 min. The reaction mixture was evaporated to dryness and the resulting crude compound was used without further purification in the next coupling reaction following the procedure described above.

**1a'**
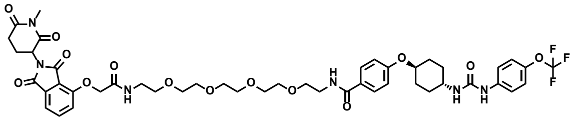
(74% yield). $^1$H NMR (400 MHz, $CDCl_3$) δ 7.81 – 7.70 (m, 3H), 7.71 (s, 2H), 7.44 – 7.35 (m, 2H), 7.23 – 7.10 (m, 4H), 6.93 – 6.86 (m, 2H), 5.01 – 4.93 (m, 1H), 4.67 (s, 2H), 4.36 (s, 4H), 4.17 (d, $J$ = 10.2 Hz, 1H), 3.70 – 3.55 (m, 15H), [PEG-H potentially overlap with a water peak], 3.23 (s, 3H), 3.05 – 2.91 (m, 1H), 2.87 – 2.72 (m, 2H), 2.17 – 2.05 (m, 5H), 1.56 (t, $J$ = 11.1 Hz, 2H), 1.30 – 1.16 (m, 2H).
$^{13}$C NMR (151 MHz, $CDCl_3$) δ 168.81, 168.46, 168.19, 166.72, 160.68, 159.58, 159.31, 156.40, 154.32, 145.58, 137.11, 136.47, 133.58, 129.22, 125.86, 123.02, 122.66, 122.14, 121.32, 119.76, 117.62, 115.37, 74.85, 70.39, 70.33, 70.31, 70.17, 70.07, 69.70, 69.30, 67.89 [70-67 ppm region potentially several PEG peaks overlap], 50.04, 48.41, 40.09, 39.20, 31.85, 30.53, 29.82, 27.32, 21.90.
$^{19}$F NMR (376 MHz, DMSO) δ -56.99.
(+) calcd for $(M+H)^+$985.3. Found 985.5. Purity (HPLC-UV 230 nm): >99%. ($^t$R= 5.64 min).

**1a**

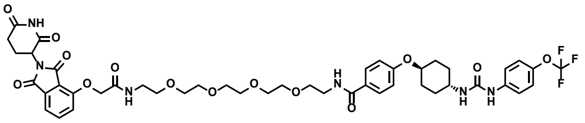

(75% yield). $^{1}$H NMR (600 MHz, DMSO) δ 11.11 (s, 1H), 8.54 (d, *J* = 5.8 Hz, 1H), 8.33 (t, *J* = 5.6 Hz, 1H), 8.00 (t, *J* = 5.7 Hz, 1H), 7.84 – 7.75 (m, 3H), 7.52 – 7.44 (m, 3H), 7.40 (d, *J* = 8.5 Hz, 1H), 7.24 – 7.19 (m, 2H), 7.02 – 6.96 (m, 2H), 6.21 (d, *J* = 7.4 Hz, 1H), 5.12 (dd, *J* = 12.9, 5.5 Hz, 1H), 4.79 (s, 2H), 4.43 (td, *J* = 9.8, 4.8 Hz, 1H), 3.57 – 3.43 (m, 15H), 3.38 (q, *J* = 6.0 Hz, 2H), 3.31 (t, *J* = 5.7 Hz, 1H), 2.89 (ddd, *J* = 17.1, 13.9, 5.5 Hz, 1H), 2.63 – 2.50 (m, 2H), 2.07 – 2.01 (m, 3H), 1.96 – 1.91 (m, 2H), 1.53 – 1.43 (m, 2H), 1.42 – 1.32 (m, 2H). $^{13}$C NMR (151 MHz, DMSO) δ 172.76, 169.86, 166.88, 166.73, 165.72, 165.44, 159.66, 154.98, 154.40, 141.94, 139.81, 136.93, 133.04, 129.00, 126.40, 121.62, 121.05, 120.34, 118.53, 116.76, 116.04, 114.83, 74.11, 69.77, 69.71, 69.61, 69.59, 69.00, 68.80, 67.50, [70-67 ppm region several PEG peaks overlap], 48.80, 47.17, 40.06, 39.94, 39.80, 39.66, 39.52, 39.38, 39.24, 39.10, 39.08, 38.40, 30.94, 29.98, 29.66, 21.99.
$^{19}$F NMR (376 MHz, DMSO) δ -56.93.
(+) calcd for $(M+H)^{+}$971.4. Found 971.3. Purity (HPLC-UV 230 nm): >99%. ($^{t}$R= 5.42 min).

**1b**

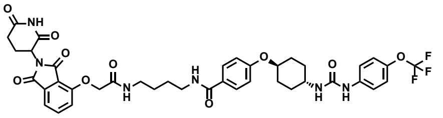

(84% yield). $^{1}$H NMR (600 MHz, DMSO) δ 11.11 (s, 1H), 8.53 (s, 1H), 8.28 (t, *J* = 5.7 Hz, 1H), 7.98 (t, *J* = 5.8 Hz, 1H), 7.83 – 7.75 (m, 3H), 7.50 – 7.44 (m, 3H), 7.39 (d, *J* = 8.5 Hz, 1H), 7.24 – 7.19 (m, 2H), 7.02 – 6.96 (m, 2H), 6.21 (d, *J* = 7.6 Hz, 1H), 5.12 (dd, *J* = 12.8, 5.5 Hz, 1H), 4.77 (s, 2H), 4.46 – 4.40 (m, 1H), 3.57 – 3.49 (m, 1H), 3.23 (q, *J* = 6.4 Hz, 2H), 3.18 (q, *J* = 6.4 Hz, 2H), 2.96 – 2.84 (m, 1H), 2.62 – 2.52 (m, 2H), 2.08 – 1.98 (m, 3H), 1.96 – 1.91 (m, 2H), 1.51 – 1.45 (m, 6H),1.42 – 1.33 (m, 2H).
$^{13}$C NMR (151 MHz, DMSO) δ 172.76, 169.87, 166.68, 165.56, 165.48, 159.56, 155.08, 154.39, 141.94, 139.80, 136.92, 133.03, 128.94, 126.65, 121.62, 121.03, 120.38, 119.35, 118.53, 116.80, 116.00, 114.82, 74.09, 67.63, 48.79, 47.17, 38.73, 38.16, 30.93, 29.97, 29.66, 26.66, 26.62, 21.98. $^{19}$F NMR (376 MHz, DMSO) δ -58.62.
(+) calcd for $(M+H)^{+}$823.3. Found 823.3. Purity (HPLC-UV 230 nm): >99%. ($^{t}$R= 5.49 min).

**1c**

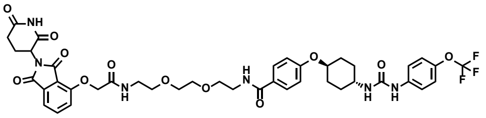

(49% yield). $^{1}$H NMR (400 MHz, $CDCl_3$) δ 8.81 (s, 1H), 7.73 (dd, *J* = 8.5, 7.0 Hz, 3H), 7.68 (s, 1H), 7.53 (d, *J* = 7.3 Hz, 1H), 7.37 – 7.31 (m, 3H), 7.16 (d, *J* = 8.5 Hz, 3H), 7.06 (s, 1H), 6.85 (d, *J* = 8.8 Hz, 2H), 4.94 (dd, *J* = 12.0, 5.9 Hz, 1H), 4.62 (s, 2H), 4.17 (s, 1H), 3.78 – 3.48 (m, 13H), 2.91 – 2.60 (m, 4H), 2.20 – 2.01 (m, 5H), 1.57 (q, *J* = 11.3 Hz, 2H), 1.32 – 1.14 (m, 2H).
$^{13}$C NMR (151 MHz, $CDCl_3$) δ 171.43, 168.53, 168.00, 167.42, 166.69, 166.17, 160.69, 155.60, 145.14, 137.32, 137.27, 133.70, 129.19, 126.47, 122.17, 121.80, 121.48, 119.78, 119.65, 118.13, 117.63, 115.46, 75.19, 70.23, 70.10, 69.93, 69.65, 68.03, 49.40, 48.34, 40.08, 39.26, 31.45, 30.84, 30.11, 22.84.$^{19}$F NMR (376 MHz, DMSO) δ -56.93.
(+) calcd for $(M+H)^{+}$883.3. Found 883.3. Purity (HPLC-UV 230 nm): >99%. ($^{t}$R= 5.46 min).

**1d**

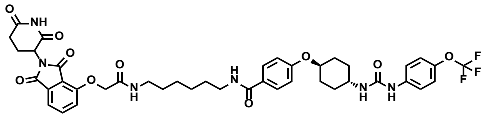

(80% yield). $^{1}$H NMR (600 MHz, DMSO) δ 11.10 (s, 1H), 8.52 (s, 1H), 8.24 (t, *J* = 5.6 Hz, 1H), 7.93 (t, *J* = 5.8 Hz, 1H), 7.80 (dd, *J* = 8.5, 7.3 Hz, 1H), 7.79 – 7.74 (m, 2H), 7.50 – 7.43 (m, 3H), 7.38 (d, *J* = 8.5 Hz, 1H), 7.20 (d, *J* = 8.7 Hz, 2H), 7.00 – 6.94 (m, 2H), 6.20 (d, *J* = 7.6 Hz, 1H), 5.11 (dd, *J* = 12.8, 5.4 Hz, 1H), 4.76 (s, 2H), 4.41 (tt, *J* = 10.0, 4.0 Hz, 1H), 3.52 (tdt, *J* = 11.0, 7.7, 4.0 Hz, 1H), 3.20 (q, *J* = 6.6 Hz, 2H), 3.13 (q, *J* = 6.6 Hz, 2H), 2.88 (ddd, *J* = 17.1, 13.9, 5.5 Hz, 1H), 2.62 – 2.49 (m, 2H), 2.02 (dddt, *J* = 11.0, 8.0, 5.5, 3.5 Hz, 3H), 1.92 (dt, *J* = 13.4, 4.1 Hz, 2H), 1.51 – 1.40 (m, 6H), 1.36 (tdd, *J* = 12.8, 10.5, 3.2 Hz, 2H), 1.32 – 1.25 (m, 4H).

$^{13}$C NMR (151 MHz, DMSO) δ 172.77, 169.88, 166.74, 166.63, 165.54, 159.54, 155.05, 154.40, 141.95, 139.81, 136.92, 133.03, 128.94, 126.72, 121.62, 121.05, 120.37, 119.36, 118.54, 116.82, 116.03, 114.83, 74.10, 67.63, 48.80, 47.18, 39.01, 38.27, 30.94, 29.98, 29.66, 29.19, 28.98, 26.18, 26.04, 21.99. $^{19}$F NMR (376 MHz, DMSO) δ -58.59.

(+) calcd for $(M+H)^+$851.3. Found 851.3. Purity (HPLC-UV 230 nm): >99%. ($^t$R= 5.72 min).

**1e**

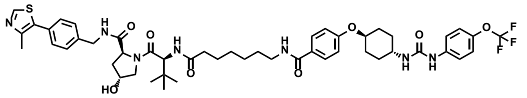

$^{1}$H NMR (600 MHz, DMSO) δ 8.99 (s, 1H), 8.56 (t, *J* = 6.1 Hz, 1H), 8.53 (s, 1H), 8.25 (t, *J* = 5.6 Hz, 1H), 7.85 (d, *J* = 9.4 Hz, 1H), 7.81 – 7.74 (m, 2H), 7.50 – 7.45 (m, 2H), 7.42 (d, *J* = 8.3 Hz, 2H), 7.40 – 7.36 (m, 2H), 7.24 – 7.19 (m, 2H), 7.01 – 6.94 (m, 2H), 6.21 (d, *J* = 7.5 Hz, 1H), 4.54 (d, *J* = 9.4 Hz, 1H), 4.46 – 4.40 (m, 3H), 4.35 (s, 1H), 4.22 (dd, *J* = 15.9, 5.5 Hz, 1H), 3.70 – 3.62 (m, 2H), 3.55 – 3.49 (m, 1H), 3.23 – 3.18 (m, 2H), 2.44 (s, 3H), 2.26 (dt, *J* = 14.6, 7.6 Hz, 1H), 2.12 (ddd, *J* = 14.2, 8.0, 6.3 Hz, 1H), 2.04 (s, 2H), 2.07 – 2.00 (m, 1H), 1.94 (d, *J* = 3.8 Hz, 1H), 1.94 – 1.86 (m, 2H), 1.55 – 1.45 (m, 6H), 1.41 – 1.32 (m, 2H), 1.30– 1.25 (m, 4H), 0.93 (s, 9H).

$^{13}$C NMR (151 MHz, DMSO) δ 172.09, 171.96, 169.72, 165.54, 159.54, 154.40, 151.48, 147.68, 141.95, 139.81, 139.52, 131.18, 129.62, 128.95, 128.64, 127.42, 126.73, 121.62, 121.05, 119.36, 118.54, 114.82, 74.09, 68.86, 58.68, 56.35, 56.28, 47.17, 41.64, 37.95, 35.19, 34.84, 29.98, 29.66, 29.15, 28.46, [31-28 ppm region alkyl peaks overlap], 26.38, 26.29, 25.42, 15.93.

$^{19}$F NMR (376 MHz, DMSO) δ -56.94.

(+) calcd for $(M+H)^+$978.4. Found 978.3. Purity (HPLC-UV 230 nm): >99%. ($^t$R= 5.65 min).

**1f**

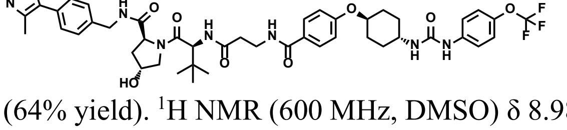

(64% yield). $^{1}$H NMR (600 MHz, DMSO) δ 8.98 (s, 1H), 8.57 (t, *J* = 6.1 Hz, 1H), 8.52 (s, 1H), 8.25 (t, *J* = 5.5 Hz, 1H), 7.99 (d, *J* = 9.3 Hz, 1H), 7.80 – 7.74 (m, 2H), 7.50 – 7.44 (m, 2H), 7.42 (d, *J* = 8.2 Hz, 2H), 7.38 (d, *J* = 8.3 Hz, 2H), 7.22 (d, *J* = 8.6 Hz, 2H), 7.01 – 6.96 (m, 2H), 6.52 (s, 1H), 6.20 (d, *J* = 7.6 Hz, 1H), 5.14 (s, 1H), 4.56 (d, *J* = 9.3 Hz, 1H), 4.47 – 4.39 (m, 2H), 4.36 (s, 1H), 4.22 (dd, *J* = 15.9, 5.5 Hz, 1H), 3.71 – 3.63 (m, 2H), 3.52 (dq, *J* = 10.6, 5.4 Hz, 1H), 3.43 (s, 1H), 2.51 – 2.41 (m, 2H), 2.44 (s, 3H), 2.07 – 2.01 (m, 3H), 1.92 (tt, *J* = 12.7, 6.6 Hz, 3H), 1.52 – 1.43 (m, 2H), 1.41 – 1.32 (m, 2H), 0.92 (s, 9H).

$^{13}$C NMR (151 MHz, DMSO) δ 172.40, 170.90, 170.07, 166.09, 160.10, 154.87, 151.93, 148.19, 142.42, 140.28, 139.97, 131.64, 130.11, 129.43, 129.12, 127.89, 127.00, 122.10, 121.52, 119.01, 115.31, 74.58, 69.37, 59.19, 56.94, 56.84, 47.65, 42.12, 38.42, 36.63, 35.71, 35.44, 30.46, 30.14, 26.84, 16.42. $^{19}$F NMR (376 MHz, DMSO) δ -56.96.

(+) calcd for $(M+H)^+$922.4. Found 922.3. Purity (HPLC-UV 230 nm): >99%. ($^t$R= 5.42 min).

**1g**

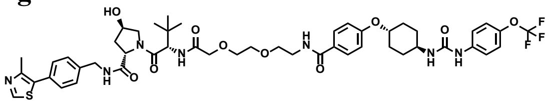

(91% yield). $^{1}$H NMR (600 MHz, DMSO) δ 8.96 (s, 1H), 8.57 (t, $J$ = 6.0 Hz, 1H), 8.52 (s, 1H), 8.33 (t, $J$ = 5.7 Hz, 1H), 7.83 – 7.75 (m, 2H), 7.50 – 7.42 (m, 3H), 7.39 (s, 4H), 7.28 – 7.19 (m, 2H), 7.02 – 6.96 (m, 2H), 6.51 (s, 1H), 6.20 (d, $J$ = 7.6 Hz, 1H), 5.16 (s, 1H), 4.57 (d, $J$ = 9.6 Hz, 1H), 4.49 – 4.34 (m, 3H), 4.25 (dd, $J$ = 15.7, 5.7 Hz, 1H), 3.97 (d, $J$ = 1.6 Hz, 2H), 3.70 – 3.51 (m, 7H), 3.42 (t, $J$ = 6.0 Hz, 2H), [PEG peak overlaps with DMSO signals], 2.43 (s, 3H), 2.06 (dd, $J$ = 17.4, 11.2 Hz, 3H), 1.91 (qd, $J$ = 10.3, 5.8 Hz, 3H), 1.48 (q, $J$ = 11.1 Hz, 2H), 1.36 (q, $J$ = 11.3 Hz, 2H), 0.94 (s, 9H).
$^{13}$C NMR (151 MHz, DMSO) δ 172.17, 169.67, 169.08, 166.24, 160.12, 154.87, 151.91, 148.22, 142.43, 140.28, 139.87, 131.60, 130.19, 129.49, 129.17, 127.93, 126.88, 122.10, 121.52, 119.01, 115.32, 74.58, 70.86, 70.02, 69.82, 69.62, 69.33, 59.20, 57.08, 56.18, 47.64, 42.16, 38.41, [PEG peak overlaps with DMSO signals], 36.20, 30.45, 30.13, 26.66, 16.40.
$^{19}$F NMR (376 MHz, DMSO) δ -56.95.
(+) calcd for (M+H)$^{+}$996.4. Found 996.3. Purity (HPLC-UV 230 nm): >99%. ($^{t}$R= 5.57 min).

**2a'**

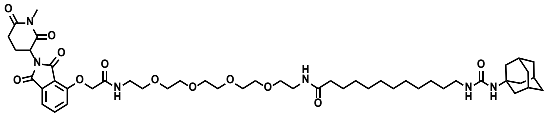

(80% yield). $^{1}$H NMR (400 MHz, CDCl$_3$) δ 7.74 (dd, $J$ = 8.4, 7.4 Hz, 1H), 7.61 (t, $J$ = 5.6 Hz, 1H), 7.54 (dd, $J$ = 7.4, 0.6 Hz, 1H), 7.20 (dd, $J$ = 8.4, 0.7 Hz, 1H), 6.39 (s, 1H), 5.01 – 4.92 (m, 1H), 4.66 (s, 2H), 3.69 – 3.49 (m, 18H), 3.42 (q, $J$ = 5.2 Hz, 2H), 3.21 (s, 3H), 3.08 – 2.93 (m, 3H), 2.87 – 2.70 (m, 2H), 2.20 – 2.14 (m, 2H), 2.14 – 2.08 (m, 1H), 2.08 – 2.03 (m, 3H), 2.00 – 1.90 (m, 6H), 1.65 (s, 6H), 1.63 – 1.55 (m, 2H), 1.54 – 1.44 (m, 2H), 1.35 – 1.20 (m, 14H).
$^{13}$C NMR (151 MHz, CDCl$_3$) δ 173.78, 171.20, 168.79, 167.25, 166.91, 166.08, 158.47, 154.62, 137.10, 133.81, 119.65, 118.24, 117.49, 70.63, 70.58, 70.41, 70.27, 70.13, 69.68, 68.19, [70-67 ppm region several PEG peaks overlap], 51.75, 50.15, 42.47, 41.06, 39.31, 39.15, 36.73, 36.41, 32.01, 29.63, 29.48, 29.45, 29.43, 29.36, 29.33, 29.25, 27.41, 26.93, 25.84, [35-25 ppm region several alkyl peaks overlap], 22.07.
(+) calcd for (M+H)$^{+}$939.5. Found 939.5. Purity (HPLC-UV 230 nm): 99%. ($^{t}$R= 6.17 min).

**2a**

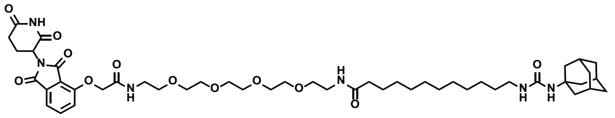

(79% yield). $^{1}$H NMR (600 MHz, DMSO) δ 11.11 (s, 1H), 8.01 (s, 1H), 7.84 – 7.78 (m, 2H), 7.50 (d, $J$ = 7.2 Hz, 1H), 7.40 (d, $J$ = 8.5 Hz, 1H), 6.52 (s, 1H), 5.57 (t, $J$ = 5.7 Hz, 1H), 5.41 (s, 1H), 5.12 (dd, $J$ = 12.9, 5.4 Hz, 1H), 4.79 (s, 2H), 3.53 – 3.44 (m, 15H), 3.37 (t, $J$ = 6.0 Hz, 2H), 3.17 (q, $J$ = 5.9 Hz, 2H), 2.89 (q, $J$ = 6.2 Hz, 3H), 2.03 (t, $J$ = 7.5 Hz, 3H), 1.97 (s, 3H), 1.84 (d, $J$ = 2.9 Hz, 6H), 1.59 (s, 6H), 1.46 (d, $J$ = 7.6 Hz, 2H), 1.30 (s, 3H), 1.23 (s, 14H).
$^{13}$C NMR (151 MHz, DMSO) δ 172.65, 172.08, 169.75, 166.78, 166.62, 165.33, 156.92, 154.88, 136.84, 132.94, 120.25, 116.66, 115.94, 69.69, 69.67, 69.62, 69.60, 69.51, 69.44, 69.05, 68.71, 67.40, 49.19, 48.70, 41.94, 38.63, 38.30, 36.03, 35.19, 30.84, 29.91, 28.93, 28.86, 28.84, 28.82, 28.70, 28.69, 28.54, 26.32, 25.15, 21.89.
(+) calcd for (M+H)$^{+}$925.5. Found 925.5. Purity (HPLC-UV 230 nm): >99%. ($^{t}$R= 5.92 min).

**2b**

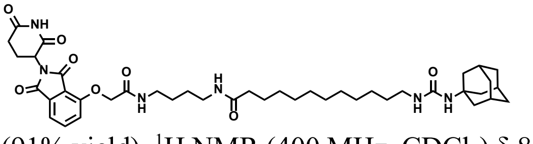

(91% yield). $^1$H NMR (400 MHz, $CDCl_3$) δ 8.99 (s, 1H), 7.75 (dd, *J* = 8.4, 7.4 Hz, 1H), 7.66 (t, *J* = 5.7 Hz, 1H), 7.57 (d, *J* = 0.6 Hz, 1H), 7.21 (dd, *J* = 8.5, 0.6 Hz, 1H), 5.78 (t, *J* = 5.7 Hz, 1H), 5.02 – 4.93 (m, 1H), 4.65 (q, *J* = 14.2 Hz, 2H), 3.42 (dt, *J* = 13.1, 6.6 Hz, 1H), 3.37 – 3.27 (m, 3H), 3.06 (t, *J* = 7.1 Hz, 2H), 2.95 – 2.84 (m, 1H), 2.84 – 2.70 (m, 2H), 2.21 – 2.12 (m, 3H), 2.09 – 2.03 (m, 3H), 1.97 – 1.92 (m, 6H), 1.69 – 1.63 (m, 6H), 1.63 – 1.52 (m, 6H), 1.51 – 1.43 (m, 2H), 1.35 – 1.16 (m, 14H).
$^{13}$C NMR (151 MHz, $CDCl_3$) δ 173.73, 171.20, 168.37, 167.15, 166.68, 166.48, 158.02, 154.93, 137.29, 133.67, 120.47, 118.60, 117.85, 68.84, 51.39, 49.51, 42.59, 40.83, 39.35, 39.05, 36.95, 36.51, 31.54, 29.93, 29.69, 29.44, 29.38, 29.37, 29.29, 29.25 [1 peak around 30 ppm representing the undecane region is missing due to overlapping], 27.02, 26.93, 26.87, 25.82, 22.89.
(+) calcd for $(M+H)^+$777.5. Found 777.3. Purity (HPLC-UV 230 nm): >99%. ($^t$R= 6.01 min).

**2c**

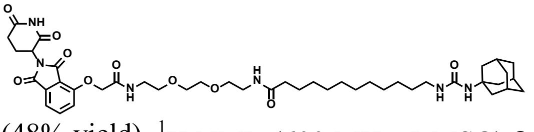

(48% yield). $^1$H NMR (600 MHz, DMSO) δ 11.11 (s, 1H), 8.01 (t, *J* = 5.7 Hz, 1H), 7.84 – 7.76 (m, 2H), 7.50 (d, *J* = 7.3 Hz, 1H), 7.40 (d, *J* = 8.6 Hz, 1H), 5.57 (t, *J* = 5.6 Hz, 1H), 5.41 (s, 1H), 5.12 (dd, *J* = 12.8, 5.5 Hz, 1H), 4.79 (s, 2H), 3.51 (h, *J* = 2.4 Hz, 4H), 3.46 (t, J = 5.7 Hz, 2H), 3.38 (t, *J* = 6.0 Hz, 2H), 3.31 (s, 1H), 3.17 (q, J = 5.9 Hz, 2H), 2.94 – 2.86 (m, 3H), 2.63 – 2.52 (m, 2H), 2.03 (dd, *J* = 8.5, 6.4 Hz, 3H), 1.97 (s, 3H), 1.83 (d, *J* = 2.9 Hz, 6H), 1.59 (d, J = 3.7 Hz, 7H), 1.46 (q, *J* = 7.3 Hz, 2H), 1.30 (d, *J* = 6.8 Hz, 1H), 1.29 (s, 2H), 1.22 (s, 16H).
$^{13}$C NMR (151 MHz, DMSO) δ 172.75, 172.21, 169.85, 166.89, 166.72, 165.43, 157.03, 154.98, 136.92, 133.04, 120.34, 116.76, 116.04, 69.57, 69.52, 69.17, 68.82, 67.50, 49.30, 48.80, 42.04, 38.73, 38.40, 36.13, 35.29, 30.94, 30.01, 29.03, 28.96, 28.94, 28.93, 28.80, 28.79, 28.65, 26.42, 25.25, 21.99.
(+) calcd for $(M+H)^+$837.5. Found 837.5. Purity (HPLC-UV 230 nm): >99%. ($^t$R= 5.92 min).

**2d**

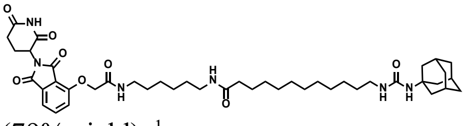

(79% yield). $^1$H NMR (600 MHz, $CDCl_3$) δ 9.24 (s, 1H), 7.74 (dd, *J* = 8.4, 7.3 Hz, 1H), 7.55 (d, *J* = 7.3 Hz, 1H), 7.52 (t, *J* = 5.7 Hz, 1H), 7.19 (d, *J* = 8.4 Hz, 1H), 5.69 (t, *J* = 6.0 Hz, 1H), 5.22 – 4.81 (m, 1H), 4.63 (dd, *J* = 14.1, 2.5 Hz, 2H), 3.42 (dq, *J* = 13.1, 6.6 Hz, 1H), 3.37 – 3.26 (m, 2H), 3.22 – 3.13 (m, 1H), 3.07 (t, *J* = 7.1 Hz, 2H), 2.96 – 2.85 (m, 1H), 2.85 – 2.74 (m, 2H), 2.22 – 2.10 (m, 3H), 2.09 – 2.03 (m, 3H), 1.97 – 1.93 (m, 6H), 1.67 – 1.63 (m, 6H), 1.60 (p, *J* = 7.0 Hz, 4H), 1.50 (p, *J* = 7.3 Hz, 2H), 1.47 – 1.39 (m, 4H), 1.39 – 1.32 (m, 2H), 1.30 – 1.20 (m, 14H).
$^{13}$C NMR (151 MHz, $CDCl_3$) δ 173.67, 171.45, 168.34, 166.86, 166.79, 166.22, 157.54, 154.64, 137.18, 133.75, 119.75, 118.38, 117.58, 68.22, 51.00, 49.49, 42.69, 40.55, 39.57, 39.19, 37.01, 36.59, 31.63, 30.27, 29.86, 29.72, 29.46, 29.40, 29.39, 29.29, 29.28, 29.26, 29.20, 26.97, 26.77, 26.56, 25.85, 22.86.
(+) calcd for $(M+H)^+$805.5. Found 805.5. Purity (HPLC-UV 230 nm): >99%. ($^t$R= 6.22 min).

**2e**

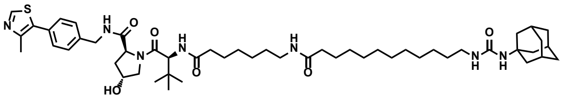

(40% yield). $^{1}$H NMR (600 MHz, DMSO) δ 9.01 (d, *J* = 2.5 Hz, 1H), 8.56 (t, *J* = 6.1 Hz, 1H), 7.83 (d, *J* = 9.4 Hz, 1H), 7.70 (t, *J* = 5.7 Hz, 1H), 7.39 (dd, *J* = 22.8, 8.2 Hz, 4H), 4.53 (d, *J* = 9.4 Hz, 1H), 4.46 – 4.39 (m, 2H), 4.36 – 4.32 (m, 1H), 4.21 (dd, *J* = 15.9, 5.5 Hz, 1H), 3.68 – 3.60 (m, 2H), 2.98 (q, *J* = 6.5 Hz, 2H), 2.88 (t, *J* = 6.9 Hz, 2H), 2.43 (s, 3H), 2.24 (dt, *J* = 14.7, 7.6 Hz, 1H), 2.09 (ddd, *J* = 14.2, 8.1, 6.3 Hz, 1H), 2.00 (t, *J* = 7.4 Hz, 3H), 1.98 – 1.94 (m, 4H), 1.89 (td, *J* = 8.4, 4.3 Hz, 1H), 1.84 – 1.80 (m, 6H), 1.62 – 1.53 (m, 6H), 1.52 – 1.39 (m, 4H), 1.38 – 1.26 (m, 4H), 1.25 – 1.16 (m, 18H), 0.95 – 0.89 (m, 9H). $^{13}$C NMR (151 MHz, DMSO) δ 172.13, 172.01, 171.96, 169.75, 157.11, 151.66, 147.46, 139.62, 131.36, 129.54, 128.67, 127.46, 68.90, 58.72, 56.38, 56.32, 49.36, 42.08, 41.68, 40.06, 38.79, 38.40, 37.98, 36.17, 35.44, 35.23, 34.87, 30.04, 29.10, 29.06, 28.99, 28.97, 28.83, 28.81, 28.69, 28.42 [One peak is overlapping around 29 ppm], 26.45, 26.40, 26.20, 25.44, 25.37, 15.84.
(+) calcd for $(M+H)^{+}$932.6. Found 932.5. Purity (HPLC-UV 230 nm): >99%. ($^{t}$R= 6.29 min).

**2f**

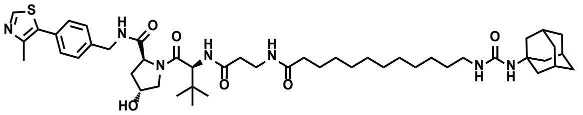

(88% yield). $^{1}$H NMR (400 MHz, $CDCl_3$) δ 8.71 (s, 1H), 7.41 – 7.30 (m, 5H), 6.64 (d, *J* = 8.5 Hz, 1H), 6.39 (t, *J* = 6.2 Hz, 1H), 4.73 (t, *J* = 8.1 Hz, 1H), 4.62 – 4.47 (m, 3H), 4.34 (dd, *J* = 15.0, 5.2 Hz, 1H), 4.06 (d, *J* = 11.3 Hz, 1H), 3.62 (dd, *J* = 11.3, 3.5 Hz, 1H), 3.46 (ddt, *J* = 36.7, 13.8, 6.9 Hz, 2H), 3.06 (t, *J* = 7.1 Hz, 2H), 2.56 – 2.39 (m, 5H), 2.40 – 2.28 (m, 1H), 2.23 – 2.07 (m, 2H), 2.08 – 2.02 (m, 3H), 1.97 – 1.90 (m, 6H), 1.70 – 1.62 (m, 6H), 1.60 – 1.51 (m, 2H), 1.50 – 1.42 (m, 2H), 1.31 – 1.19 (m, 14H), 0.94 (s, 9H).
$^{13}$C NMR (151 MHz, DMSO) δ 171.92, 171.83, 170.23, 169.49, 156.93, 151.34, 147.61, 139.40, 131.06, 129.53, 128.53, 127.31, 68.78, 58.61, 56.35, 56.23, 49.20, 41.95, 41.55, 38.64, 37.85, 36.04, 35.28, 34.93, 29.92, 28.95, 28.88, 28.83, 28.74, 28.70, 28.59, 26.33, 26.27, 25.15, 15.84.
(+) calcd for $(M+H)^{+}$876.5. Found 876.6. Purity (HPLC-UV 230 nm): >99%. ($^{t}$R= 6.02 min).

**2g**

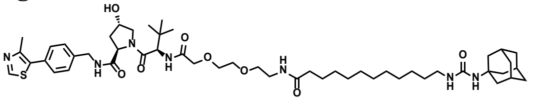

(93% yield).
$^{1}$H NMR (600 MHz, $CDCl_3$) δ 7.47 (d, *J* = 9.2 Hz, 1H), 7.37 (d, *J* = 8.2 Hz, 2H), 7.33 (d, *J* = 8.2 Hz, 2H), 7.11 (t, *J* = 5.9 Hz, 1H), 6.70 (t, *J* = 5.6 Hz, 1H), 4.68 (t, *J* = 8.0 Hz, 1H), 4.62 (d, *J* = 9.3 Hz, 1H), 4.53 (m, 2H), 4.39 (dd, *J* = 5.5, 14.9 Hz, 1H), 4.08 (d, *J* = 16.0 Hz, 1H), 3.94 (d, *J* = 16.0 Hz, 1H), 3.89 (d, *J* = 11.3 Hz, 1H), 3.73 (m, 1H), 3.64 (m, 4H), 3.54 (m, 2H), 3.43 (m, 4H), 3.06 (t, *J* = 7.1 Hz, 2H), 2.52 (s, 3H), 2.49 (q, *J* = 4.3 Hz, 1H), 2.24 (m, 2H), 2.15 (q, *J* = 7.2 Hz, 1H), 2.06 (s, 3H), 1.94 (d, *J* = 2.5 Hz, 6H), 1.65 (s, 6H), 1.61 (m, 2H), 1.45 (t, *J* = 6.8 Hz, 2H), 1.26 (t, *J* = 26.6 Hz, 16H), 0.95 (s, 9H).
$^{13}$C NMR (151 MHz, DMSO) δ 172.23, 171.68, 169.17, 168.62, 157.03, 151.43, 147.74, 139.39, 131.12, 129.71, 128.69, 127.44, 70.38, 69.53, 69.34, 69.33, 68.83, 58.72, 56.59, 55.68, 49.30, 42.04, 41.68, [1 peak potentially overlap with DMSO signal], 38.74, 38.39, 37.94, 36.13, 35.74, 35.27, 30.01, 29.04, 28.97, 28.94, 28.93, 28.83, 28.79, 28.64, 26.42, 26.27, 26.19, 25.25, 15.92.
(+) calcd for $(M+H)^{+}$950.6. Found 950.5. Purity (HPLC-UV 230 nm): 99%. ($^{t}$R= 6.11 min).
