## Supplementary material for "PROTAC-mediated selective degradation of cytosolic soluble epoxide hydrolase enhances ER-stress reduction": Original western data

**Fig. 1**

Fig. 1A Upper panel

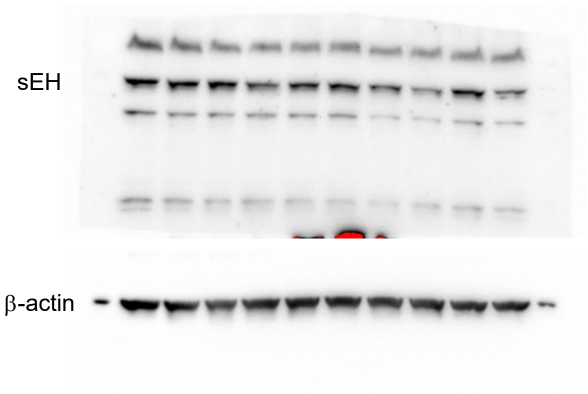


Fig. 1A Lower panel

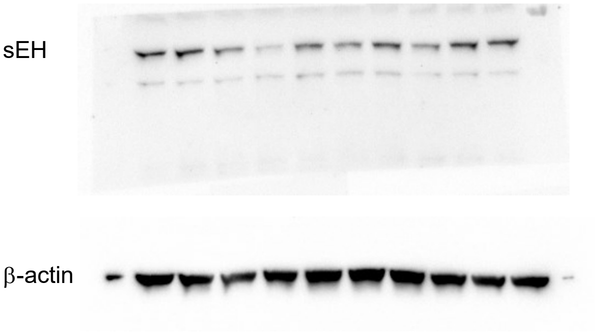


Fig. 1C

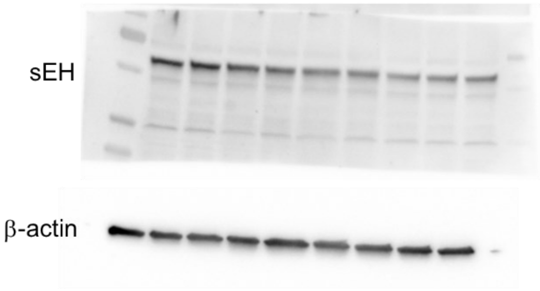


Fig. 1D  Left panel

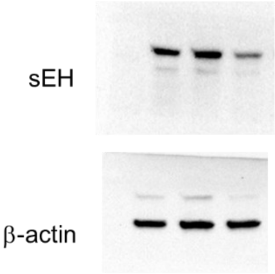


Fig. 1D  Right panel

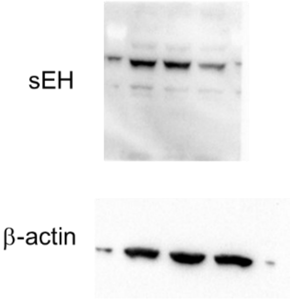

**Fig.2**

Fig. 2A

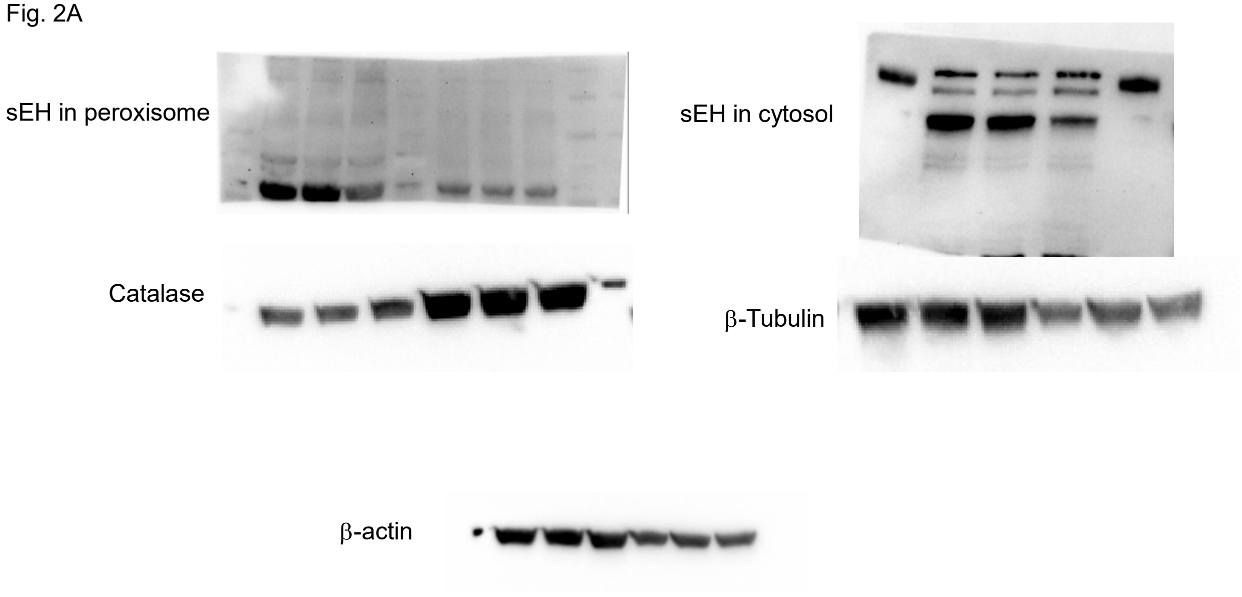

**Fig. 3**

Fig.3A Left panel

sEH

β-actin

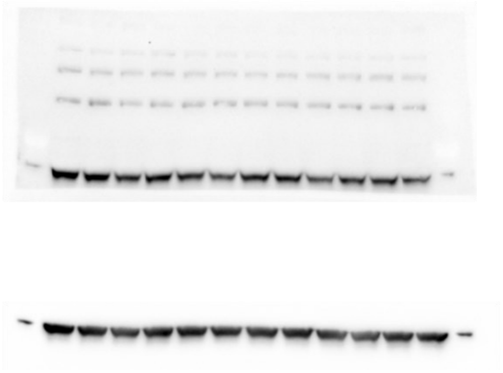

Fig.3A Right panel

sEH

β-actin

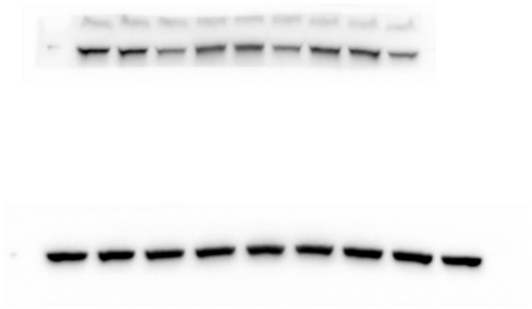

Fig.3B

sEH

β-actin

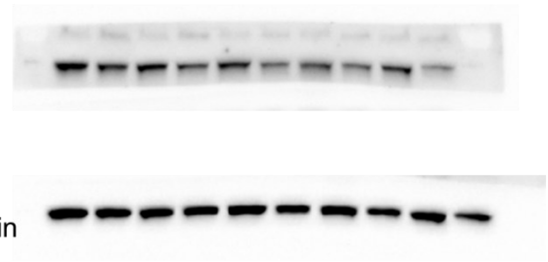

**Fig. 4**

Fig. 4A

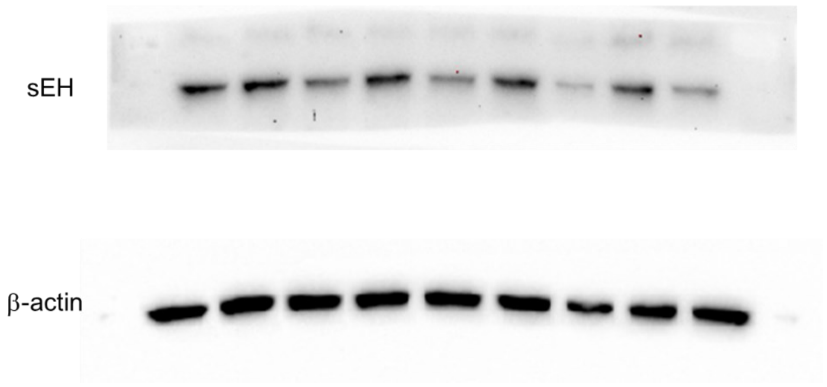


Fig. 4B

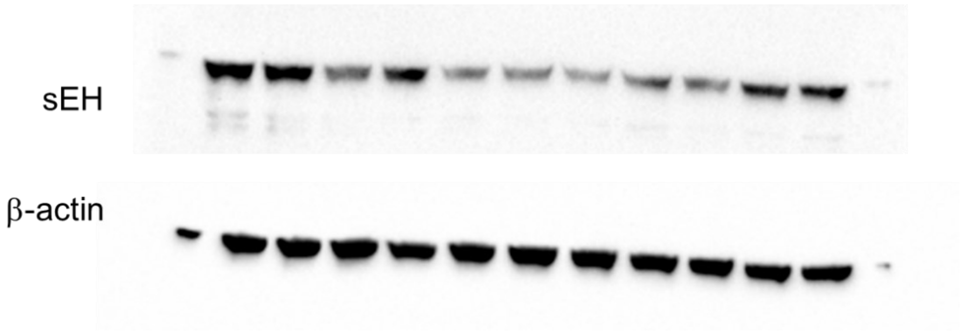

**Fig. 5**

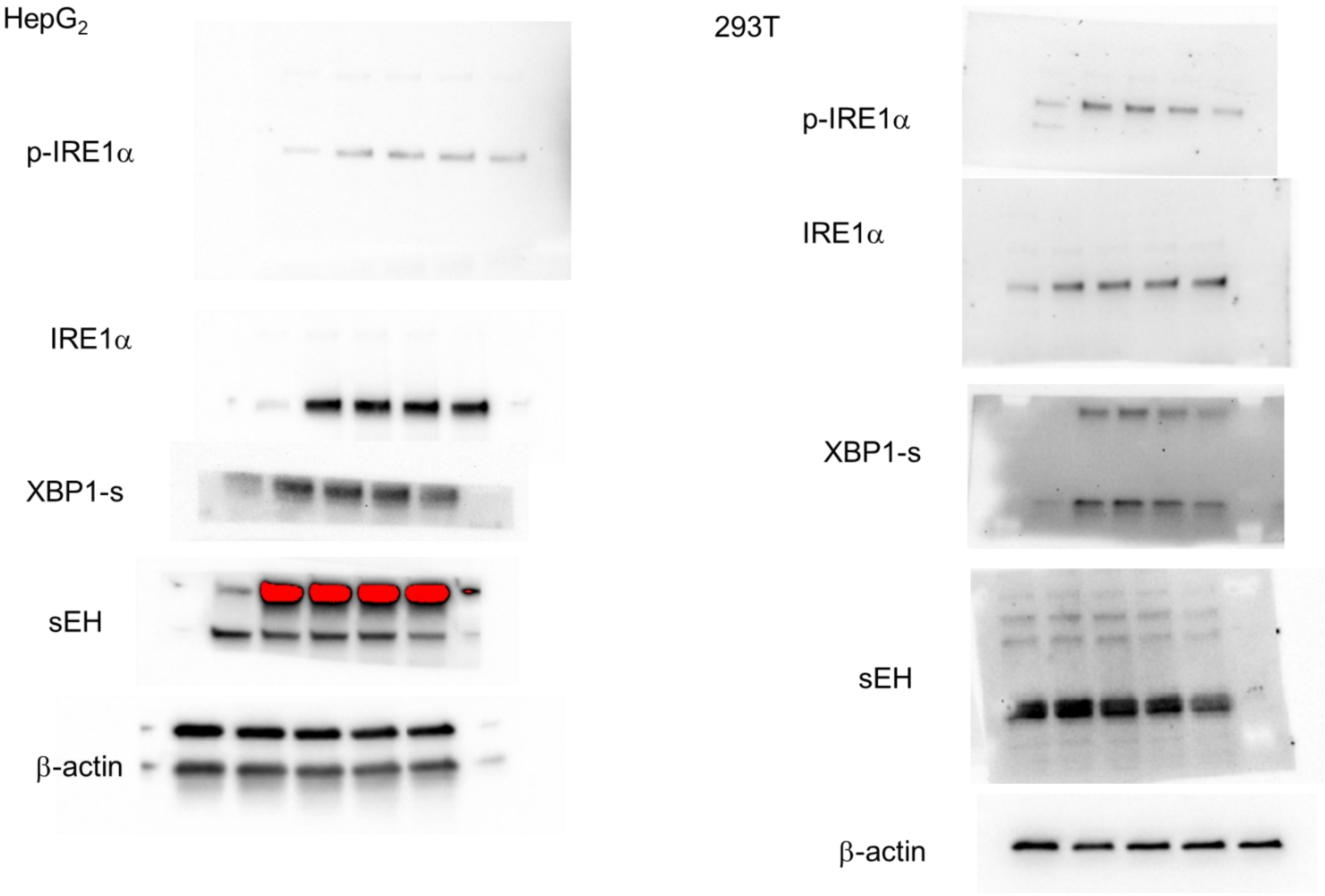
HepG$_2$
p-IRE1α
IRE1α
XBP1-s
sEH
β-actin
293T
p-IRE1α
IRE1α
XBP1-s
sEH
β-actin

**Fig. S1**

Figure S1

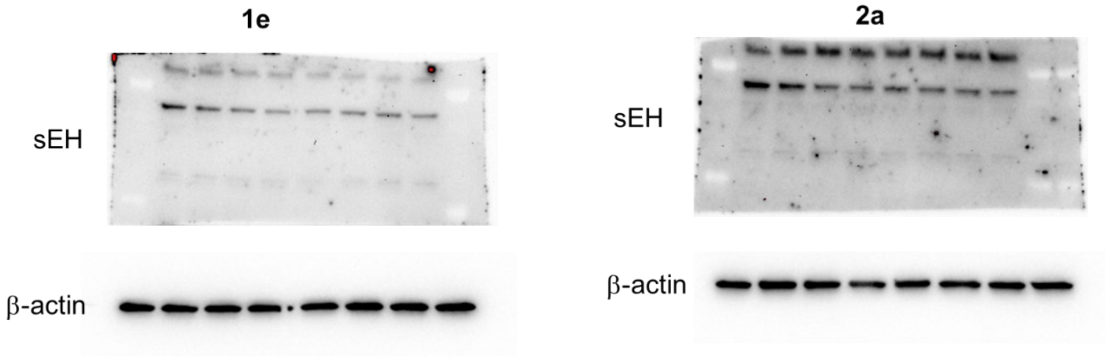

**Fig. S2**

Figure S2

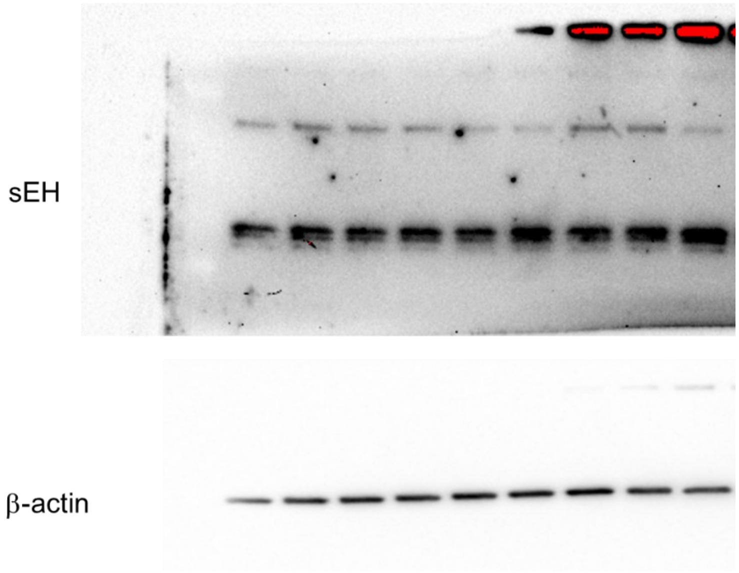

**Fig. S3**

Figure S3

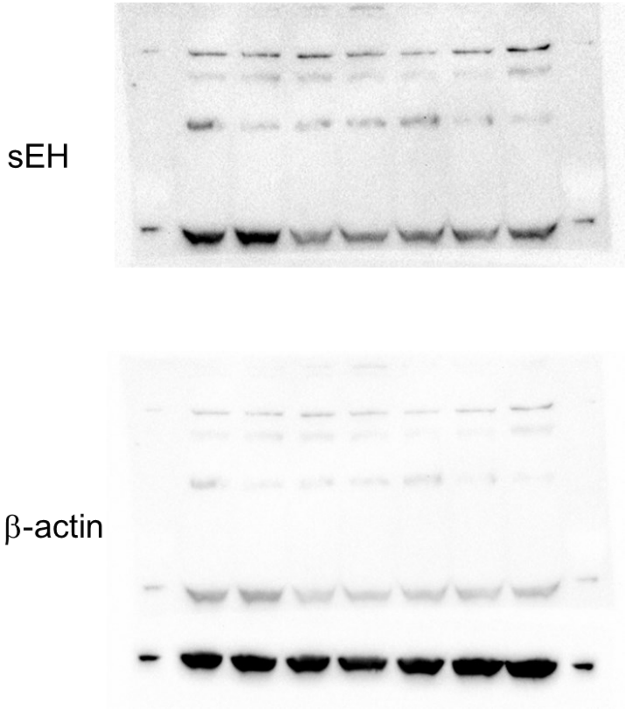


**Fig. S4**

Figure S4

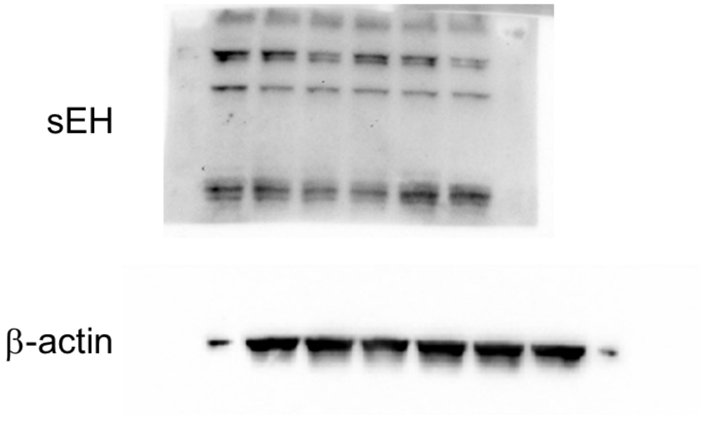

**Fig. S5**

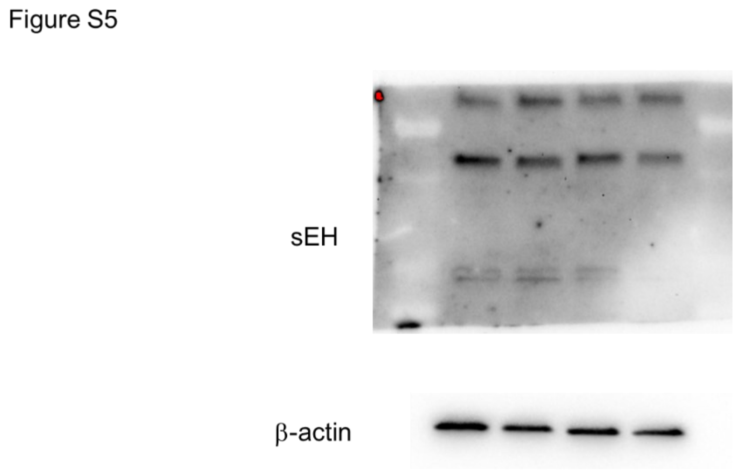
Figure S5
sEH
β-actin

**Fig. S6**

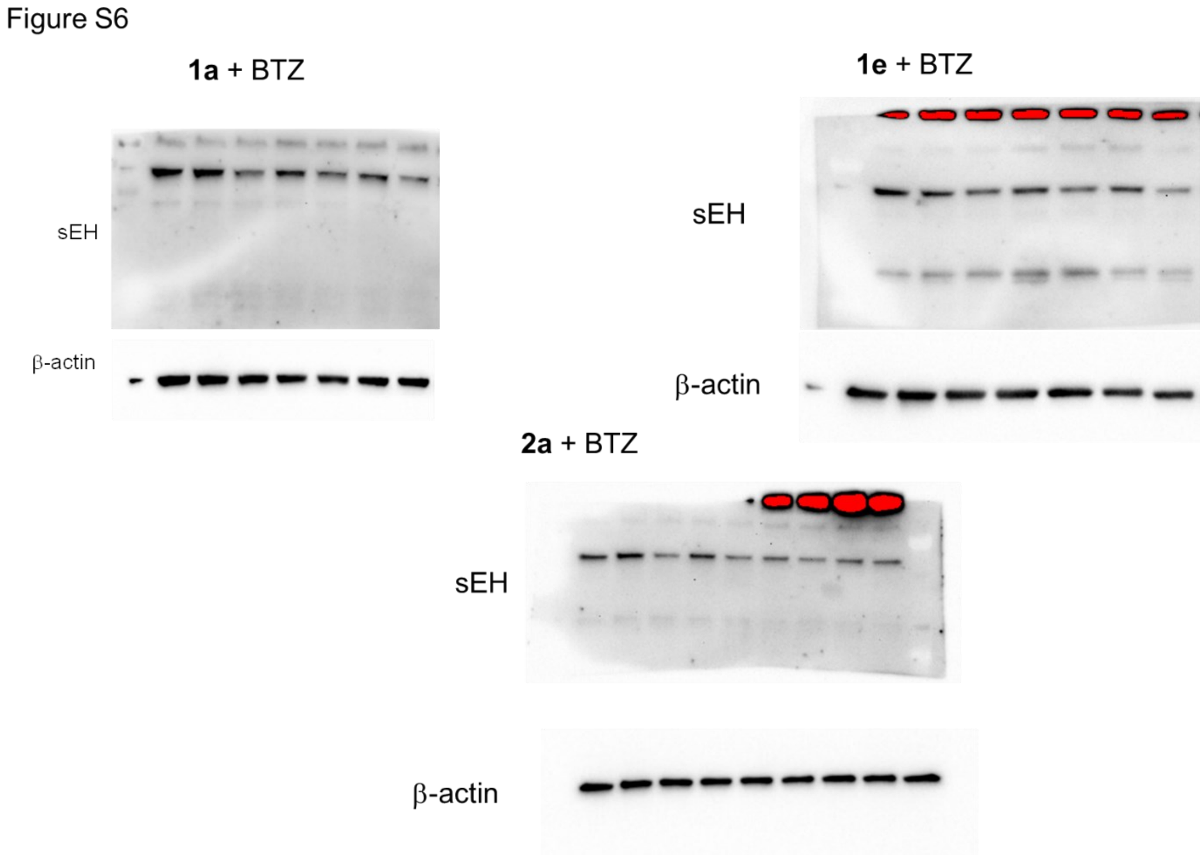
Figure S6
1a + BTZ
sEH
β-actin
1e + BTZ
sEH
β-actin
2a + BTZ
sEH
β-actin

**Fig. S7**

Figure S7

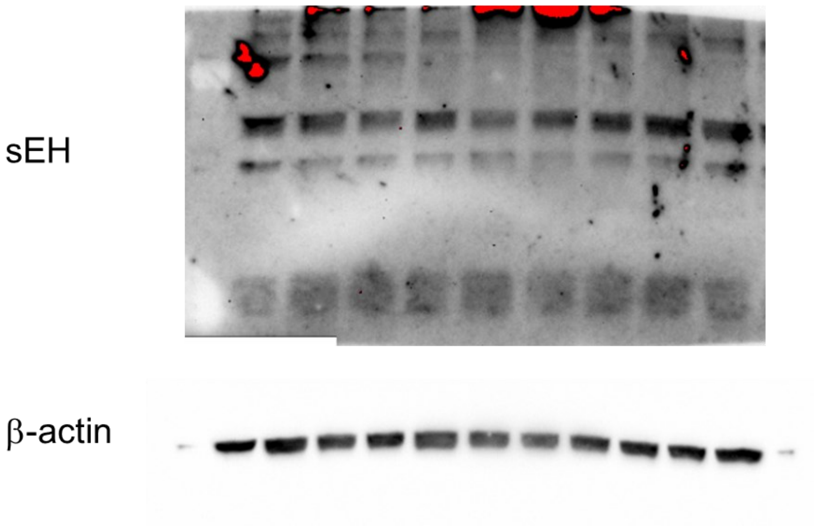


**Fig. S8**

Figure S8A

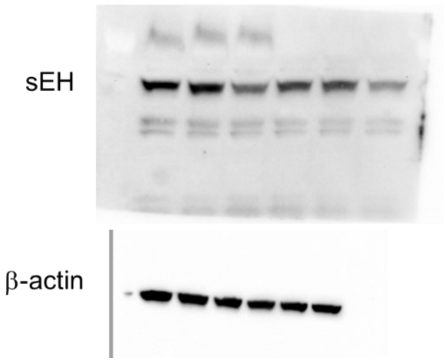

Figure S8B

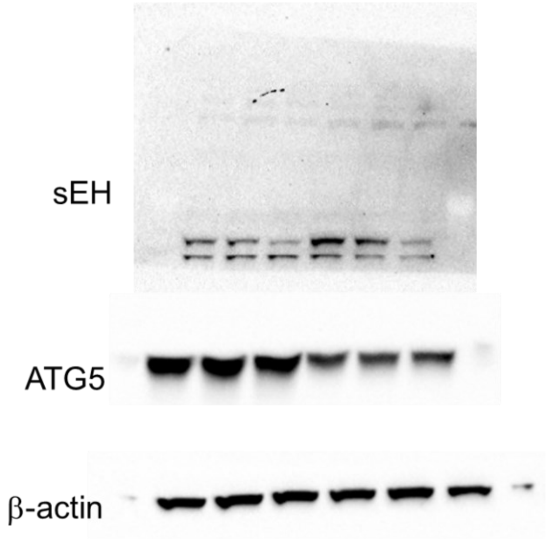


Figure S8C

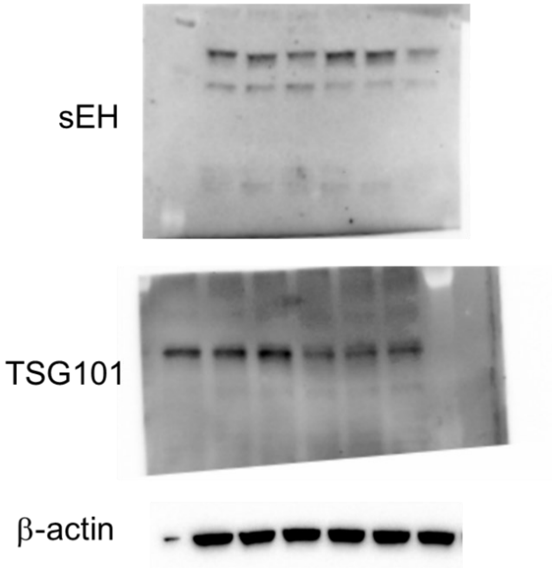

Figure S8D

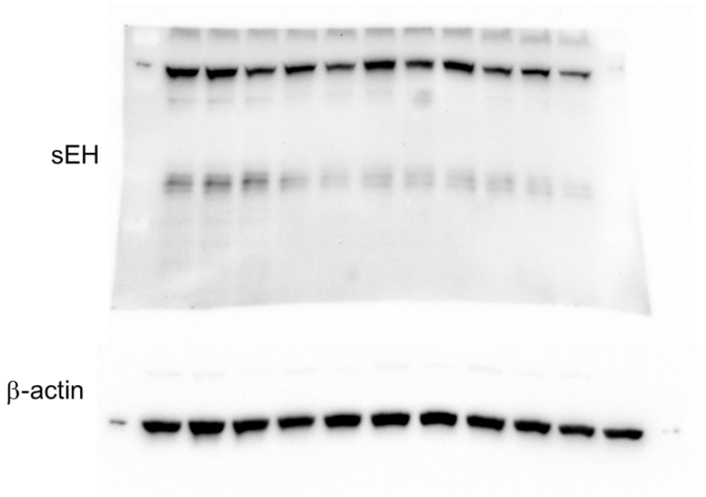
